# APOBEC3 Activation Drives Epitranscriptomic Remodeling and Clonal Hematopoiesis

**DOI:** 10.64898/2026.11.20.768132

**Authors:** Inge van der Werf, Jane Isquith, Emma Klacking, Jessica Pham, Claire Engstrom, Wenxue Ma, Shuvro P. Nandi, Rongjie Wu, Neha Katragadda, Kendale Wirtjes, Anna A. Khachatrian, Thomas Whisenant, Ludmil B. Alexandrov, Catriona H. M. Jamieson

## Abstract

Stress and inflammatory cytokine-inducible APOBEC3 (A3) enzymes promote progression of many malignancies. Previously, we identified A3C overexpression in hematopoietic stem and progenitor cells (HSPCs) in the context of spaceflight-associated accelerated aging and precancer myeloproliferative neoplasm (MPN) stem cells. However, the role of A3 in age-associated clonal hematopoiesis (CH) and precancer initiation remains unclear. To investigate the functional hierarchy among A3 enzymes, we performed lentiviral A3B, A3C, A3D, A3F, and A3G overexpression, whole genome, whole transcriptome and single-cell RNA sequencing analyses, and functional HSPC assays. In HSPCs, A3C and A3F induced more C-to-T DNA mutations; A3B, A3C, and A3G deregulated RNA splicing; and all A3 enzymes induced C-to-U RNA edits. Moreover, A3C overexpression in HSPCs enhanced replating and myeloid lineage skewing in humanized inflammatory niche mouse models. Notably, A3C-induced editing of CH-associated transcripts distinguished MPN from normal HSPCs. Thus, future strategies focused on APOBEC3C inhibition may prevent CH and precancer initiation.

## Main

Stress and inflammatory cytokine-responsive APOBEC3 (apolipoprotein B mRNA editing catalytic polypeptide-like; A3) enzymes were initially discovered as cytosine-to-uracil (C-to-U) RNA deaminases that evolved to safeguard genomic integrity by restricting both human endogenous retrovirus (HERV) retrotransposition and retroviral integration.^1^ Following DNA replication, uracil edits are read as thymine bases, resulting in cytosine-to-thymine (C-to-T) single base substitutions (SBS).^2^ However, deregulation of APOBEC3-mediated C-to-T DNA mutagenesis has emerged as a major source of mutagenesis in a broad array of cancers.^3–10^ Despite the prominent role of APOBEC3 base deaminases in cancer progression, their importance in inflammation-associated hematopoietic stem and progenitor cell (HSPC) aging, clonal hematopoiesis (CH), and precancer initiation remains undefined.^11^

Among the seven human APOBEC3 family members, APOBEC3A and APOBEC3B have been widely studied due to their elevated expression and mutagenic activity in diverse cancer types.^3–8,12,13^ Their activity generates distinctive mutational signatures that contribute to tumor heterogeneity^14^, immune evasion^15^, and therapeutic resistance.^16^ Beyond the role of APOBEC3C in DNA replication stress resistance in some cancers^17^, APOBEC3C remains less studied as a functional deaminase, although it is frequently expressed in normal tissues and exhibits higher average expression in some tumor types than APOBEC3A or APOBEC3B.^18–21^ These observations raise the possibility that APOBEC3C contributes to mechanisms distinct from, or extending beyond, those established for APOBEC3A and APOBEC3B.

During aging, HSPCs undergo a progressive decline in regenerative capacity, exhibit skewed differentiation toward the myeloid lineage, show increased sensitivity to stressors, and accumulate somatic mutations, a process known as CH, which increases susceptibility to cardiovascular mortality, myeloproliferative neoplasms (MPNs), myelodysplastic syndromes (MDS), and progression to secondary acute myeloid leukemia (sAML).^22–28^ Also, age-associated CH contributes to inflammation in the tumor microenvironment, thereby promoting solid tumor metastases^29,30^. Additional hallmarks of aging include genomic instability, upregulation of inflammatory cytokines, deregulated RNA splicing, and derepression of human endogenous retroviruses (HERVs).^11,31–34^ Our recent studies revealed increased expression of APOBEC3C in hematopoietic stem cells (HSCs) derived from patients diagnosed with MPN, and following stress-associated HSPC aging, including inflammation or microgravity in HSPC nanobioreactors and astronauts.^35–37^ This increased expression was accompanied by mutational profiles that conformed with an APOBEC3C-driven SBS mutagenesis signatures in HSPCs and mutations in CH-associated genes. However, these studies did not establish a causal role for APOBEC3C in CH or precancer initiation.

By performing APOBEC3 family member lentiviral overexpression, whole genome sequencing (WGS), whole transcriptome RNA sequencing (RNA-seq), single-cell RNA sequencing (scRNA-seq), and functional HSPC analyses, we discovered that APOBEC3 enzymes varied in their capacity to induce DNA mutations in HSPCs, but uniformly induced C-to-U RNA edits and perturbed RNA editing and splicing. Furthermore, we demonstrated that APOBEC3C induced substantially greater transcriptomic effects in aged bone marrow (ABM) HSPCs compared to cord blood (CB) HSPCs, resulting in a ten-fold increase in the number of significantly differentially expressed genes, including increased expression of RNA splicing factors, including U2AF1, ZRSR2, and SF3B2, which are associated with aberrant RNA splicing and CH, MPN, and MDS pathogenesis,^38^ alongside transcriptome-wide changes in alternative splicing characterized by exon skipping. In ABM HSPCs, genomic and epitranscriptomic (post-transcriptional) changes were accompanied by enhanced replating and myeloid skewing in humanized inflammatory niche mouse models. Furthermore, increased C-to-U RNA edits in MPN HSCs and HPCs compared to ABM counterparts, with APOBEC3C expression showing the strongest positive association with editing burden, and C-to-U RNA edits in CH-associated genes, including CUX1, across APOBEC3C lentivirally transduced CB and ABM HSPCs, as well as MPN HSCs and HPCs. Thus, this study established a previously unrecognized role for APOBEC3 in epitranscriptomic remodeling through RNA editing and RNA splicing dysregulation, providing new insight into mechanisms that may contribute to age-associated CH and precancer initiation.

## Results

### APOBEC3-induced epitranscriptomic and genomic editing in HSPCs

While APOBEC3 family members have been shown to induce C-to-T mutations, we investigated their broader roles in DNA mutagenesis and epitranscriptomic RNA editing in the context of HSPC aging and CH. We developed lentiviral vectors driven by the EF1α promoter that enabled overexpression of APOBEC3B, A3C, A3D, A3F, and A3G in immunomagnetic bead-selected CD34^+^ CB HSPCs. Subsequently, we performed WGS (90X) and RNA-seq analyses (**Fig. 1a**). While APOBEC3A and APOBEC3H transcripts were undetectable in CB HSPCs, APOBEC3C levels appear to be the highest compared with APOBEC3B, A3D, A3F, and A3G (**Fig. 1b**).

**Figure 1:**
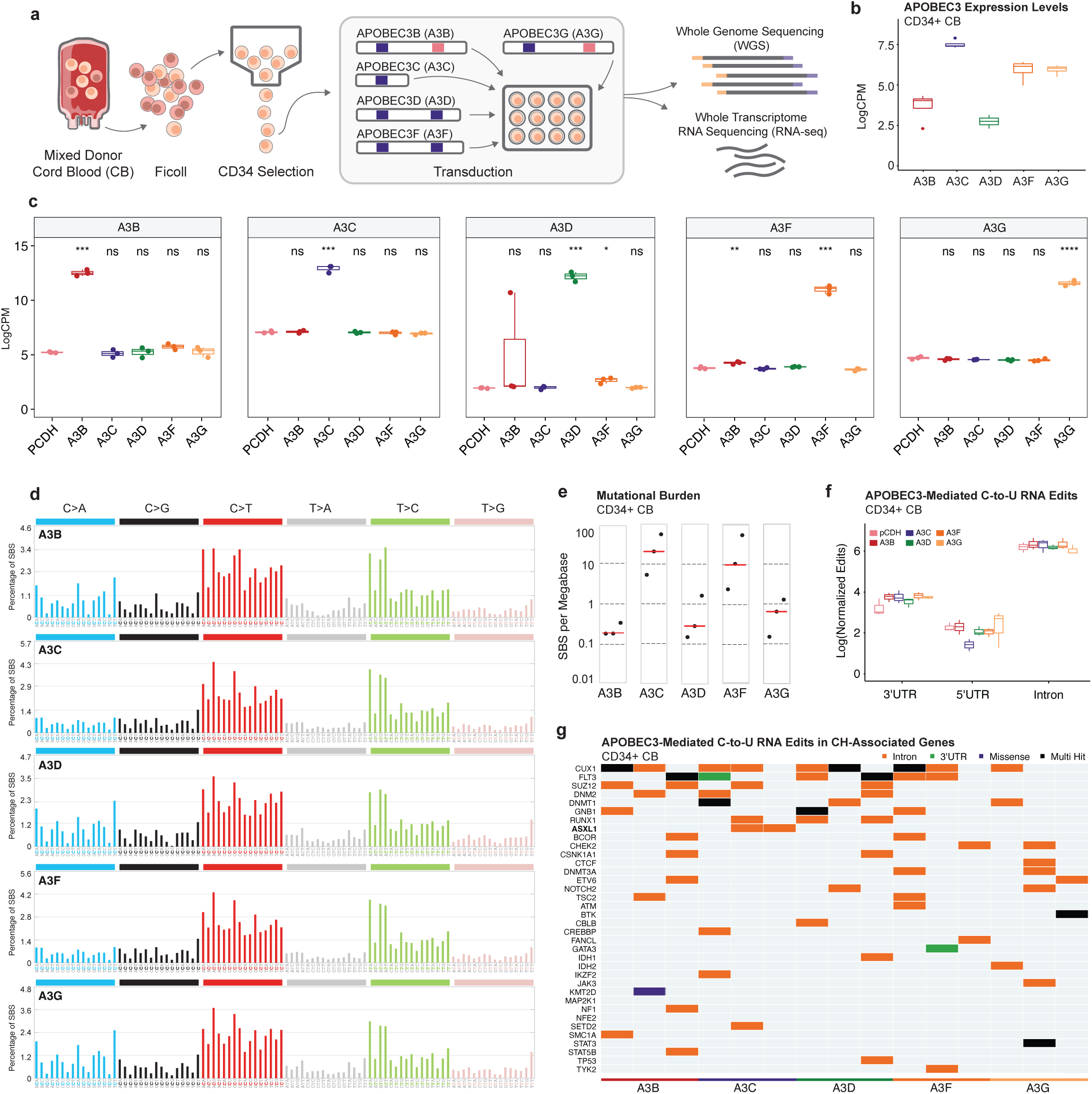
APOBEC3-induced epitranscriptomic and genomic editing in HSPCs. **a.** Schematic representation of the experimental design in cord blood (CB) hematopoietic stem and progenitor cells (HSPCs). **b.** RNA-seq based expression levels (log counts per million) of APOBEC3 A-H in CB HSPCs. **c.** RNA-seq based expression levels (log counts per million) in CB HSPCs following lentiviral transduction with pCDH, APOBEC3B, APOBEC3C, APOBEC3D, APOBEC3F, or APOBEC3G. **d.** Average mutation profiles are displayed by substitution class (C>A, C>G, C>T, T>A, T>C, T>G), with 16 bars beneath each class representing the sequence context based on the four possible nucleotides located immediately 5′ and 3′ of each mutated base. Mutations were called using Mutect2, Strelka2, Varscan2, and MuSE2, and only variants identified by at least two callers were considered. **e.** Mutational burden of single base substitutions (SBS) plotted on a logarithmic scale. Each dot represents the number of substitutions per megabase in an individual replicate; red lines indicate the median number. **f.** Boxplots of APOBEC3-mediated C-to-U RNA edits by region in CB HSPCs following lentiviral transduction with pCDH backbone, APOBEC3B, APOBEC3C, APOBEC3D, APOBEC3F, or APOBEC3G; untranslated region (UTR). **g.** Oncoplot illustrating APOBEC3-mediated C-to-U RNA edits in clonal hematopoiesis (CH)-associated genes following APOBEC3 lentiviral transduction of CB HSPCs.

First, we quantified the expression levels of each APOBEC3 family member in HSPCs following lentiviral transduction (**Fig. 1c**). Subsequently, we determined the impact of APOBEC3 lentiviral transduction on genomic integrity by identifying somatic mutations using four variant callers (Mutect2, Strelka2, Varscan2, and MuSE2), only considering mutations that were identified by two variant callers. Somatic mutations were predominantly characterized by C:G>T:A single base substitutions (SBSs) as well as T:A>C:G substitutions (**Fig. 1d; Extended Data Fig. 1a**). Mutational signature analysis revealed that the observed patterns of mutations can be recapitulated using COSMIC signature SBS1, a clock-like signature linked to cell division^39^, as well as SBS5, a clock-like signature that is independent of cell division and characteristic of quiescent cells^3,40^ (**Extended Data Fig. 1b**). Cosine similarity analysis further revealed relatively low similarity to COSMIC signatures SBS2 and SBS13, typically associated with APOBEC3A and APOBEC3B activity in yeast and solid tumors (SBS2: A3B 0.27, A3C 0.24, A3D 0.30, A3F 0.24, A3G 0.29; SBS13: A3B 0.16, A3C 0.15, A3D 0.17, A3F 0.16, A3G 0.17). In contrast, the overall mutational profiles closely recapitulated the patterns observed in MPNs derived from bulk peripheral blood (PCAWG MPN: A3B 0.91, A3C 0.92, A3D 0.91, A3F 0.92, A3G 0.91).

Interestingly, APOBEC3C- and APOBEC3F-overexpressing HSPCs exhibited a significantly higher mutational burden compared to other APOBEC3 conditions (**Fig. 1e**). Moreover, APOBEC3C- and APOBEC3F-overexpressing HSPCs exhibited double base substitutions (DBS) and insertions and deletions (InDels) with resemblance to MPN HSPCs (DBS: A3C 0.59, A3F 0.61; Indels: A3C 0.75; A3F 0.74). Further targeted analyses of 125 CH-associated genes (**Extended Data Fig. 1c**) revealed mutations in 70 genes following APOBEC3 overexpression. This included multi-hit mutations in *RUNX1*, *SETBP1*, and *JAK2*, as well as intronic mutations in CH-associated genes *DNMT3A*, *ZRSR2*, *STAT3*, *SF3A1*, and *TET2* mutations within the 5’ flank region (**Extended Data Fig. 1d**). Interestingly, *SETBP1*, *RUNX1*, and *CUX1* were mutated in all three APOBEC3C-overexpressing replicates. Recurrent and multi-hit mutations in CH-associated genes did not affect known hotspot sites and are of unknown clinical significance. Similarly, genome-wide multi-hit and missense mutations (**Extended Data Fig. 1e**) require further investigation to elucidate both the impact on HSPC function and persistence over time.

While APOBEC3 family members are widely recognized as C:G>T:A mutagenic enzymes of DNA in cancer, APOBEC3 deaminases were first described as RNA editors.^15^ To explore their role as RNA editors in HSPCs, we developed a bioinformatic pipeline to detect C-to-U editing at the transcriptome level, and did observe a trend toward increased C-to-U editing within the 3’ untranslated region (3’UTR; **Fig. 1f; Extended Data Fig. 1f, g**). Similarly, we observed trends toward reduced C-to-U editing in long non-coding RNAs (long ncRNAs), and increased editing in microRNAs (miRNAs; **Extended Data Fig. 1h**). Both require targeted validation given the low number of detected edits in these non-coding regions. Targeted analyses of CH-associated genes revealed C-to-U edits in 36 genes (**Fig. 1g**). In contrast to APOBEC-mediated DNA mutations, C-to-U RNA editing was detected across all APOBEC3 family members, with CUX1 and FLT3 consistently edited across all APOBEC3-overexpressing conditions, and ASXL1 showing editing in two out of three replicates specifically upon APOBEC3C expression (**Fig. 1g**).

### APOBEC3 activation promotes RNA splicing deregulation

Building on the observed effects of APOBEC3 on RNA editing, we investigated APOBEC3 activation in RNA rewiring through changes in gene expression and RNA splicing. Principal component analysis (PCA) demonstrated that overexpression of individual APOBEC3 family members resulted in distinct clustering of CB CD34^+^ HSPCs, indicating that each APOBEC3 enzyme induces a unique transcriptional program (**Fig. 2a**). Differential gene expression detected 577, 356, 53, 3, and 498 significantly differentially expressed genes in APOBEC3B, 3C, 3D, 3F, and 3G, respectively, compared to the pCDH backbone transduced controls (**Fig. 2b**; **Supplementary Table 1**; adjusted p-value <0.05).

**Figure 2:**
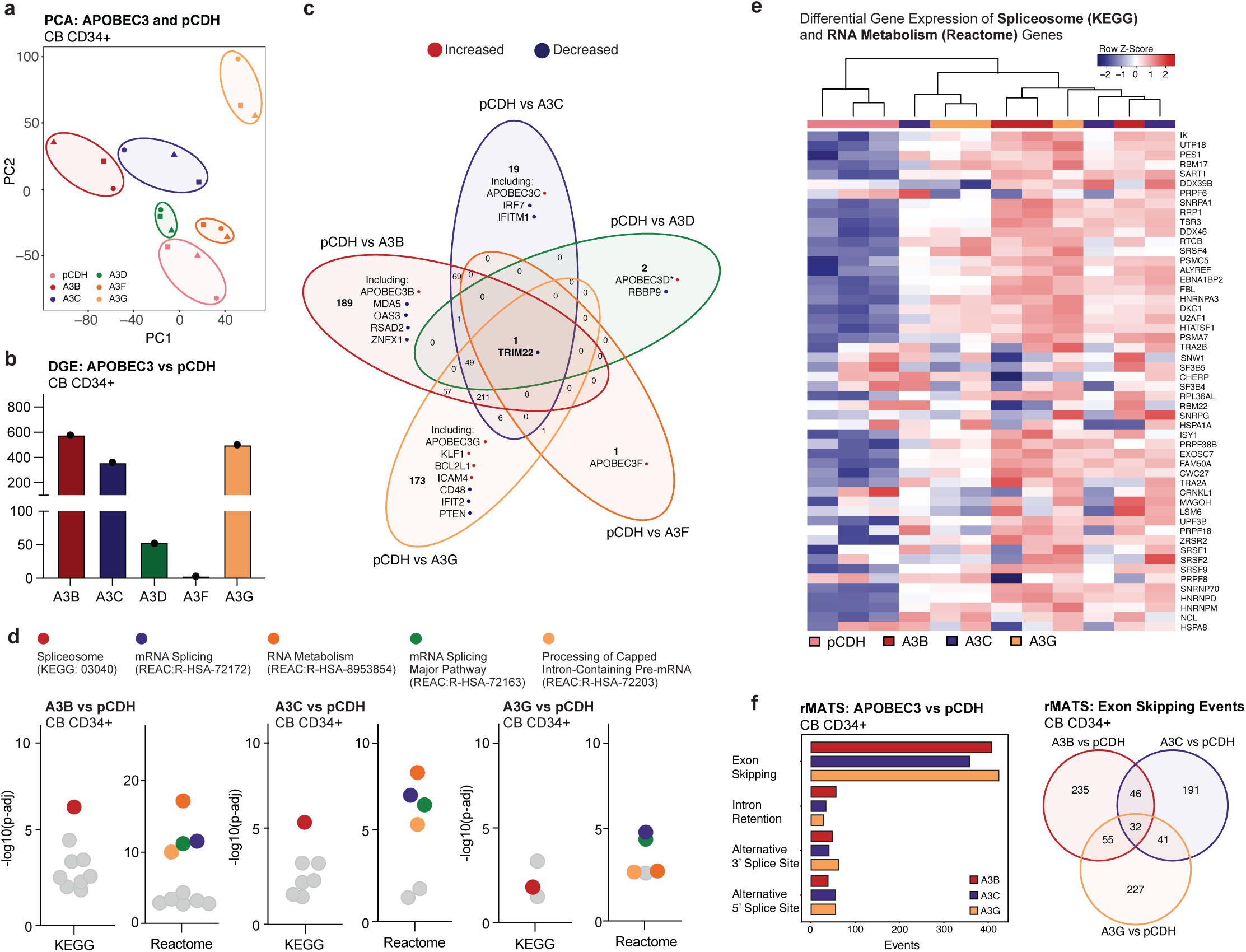
APOBEC3 activation promotes RNA splicing deregulation **a.** Principal component analysis (PCA) plot illustrating the transcriptional differences between pCDH, APOBEC3B, APOBEC3C, APOBEC3D, APOBEC3F, and APOBEC3G lentiviral transduction in cord blood (CB) hematopoietic stem and progenitor cells (HSPCs). **b.** Bar plot visualizing the absolute number of significantly differentially expressed genes induced by APOBEC3B, APOBEC3C, APOBEC3D, APOBEC3F, and APOBEC3G lentiviral transduction in CB HSPCs compared to pCDH backbone transduced controls. Genes with adjusted p-value <0.05 are included (*nominal p-value <0.001) **c.** Venn diagram visualizing the absolute number of significantly differentially expressed genes induced by APOBEC3B, APOBEC3C, APOBEC3D, APOBEC3F, and APOBEC3G lentiviral transduction in CB HSPCs compared to pCDH backbone transduced controls. Genes with adjusted p-value <0.05 are included (*nominal p-value <0.001) **d.** Enrichment analyses (gProfiler) of significantly differentially expressed genes in CB HSPCs lentivirally transduced with APOBEC3B, APOBEC3C, or APOBEC3G compared to pCDH backbone-transduced controls. Both KEGG and Reactome categories are represented. The y-axis represents the negative log of the adjusted p-value. **e.** Heatmap of gene expression levels after unsupervised hierarchical clustering of genes enriched in gProfiler analyses and part of the KEGG Spliceosome or Reactome RNA Metabolism pathways. **f.** Left: Quantification of differential splicing events, including exon skipping, intron retention, alternative 3’ splice site usage, and alternative 5’ splice site usage in CB HSPCs lentivirally transduced with APOBEC3B, APOBEC3C, or APOBEC3G, compared to pCDH backbone transduced controls, using multivariate analysis of transcript splicing (rMATS). Right: Venn diagram visualizing the absolute number of exon skipping events induced by APOBEC3B, APOBEC3C, and APOBEC3G lentiviral transduction in CB HSPCs compared to pCDH backbone transduced controls.

Initially, we evaluated the overlap between the different APOBEC3 family members and identified TRIM22, an interferon-inducible gene encoding an E3 ubiquitin ligase involved in antiviral defense, significantly downregulated across all APOBEC3 overexpression conditions (**Fig. 2c; Extended Data Fig. 2a**). Notably, TRIM22 was also found to be significantly downregulated in MPN HPCs (**Extended Data Fig. 2b**). Given the established role of TRIM22 in interferon-mediated antiviral defense, its consistent downregulation following APOBEC3 overexpression and in MPN HPCs suggests a potential link between APOBEC3 activation and altered innate immune responses in HSPCs. In this context, APOBEC3B lentiviral transduction resulted in significantly increased expression of the transduced APOBEC3B gene, alongside innate antiviral defense genes including MDA5 (IFIH1) and ZNFX1, which are involved in viral sensing and IFN signaling initiation, as well as antiviral RNA integrity regulators such as OAS3 and RSAD2 (**Fig. 2c; Extended Data Fig. 2c**); APOBEC3C lentiviral transduction uniquely and significantly induced expression of APOBEC3C, along with changes in expression of IFN response genes, IFI27 and IFITM1; and lentiviral transduction of APOBEC3G resulted in significantly differential expression of APOBEC3G, KLF1, an erythroid transcription factor essential for red blood cell maturation, the anti-apoptotic regulator BCL2L1 (BCL-XL), adhesion molecules ICAM4 and CD48, and immune-responsive IFN-related genes, such as IFIT2 and PTEN (**Fig. 2c; Extended Data Fig. 2c**).

We also observed a significant overlap of genes between lentiviral transduction of APOBEC3B, 3C and 3G (211 genes), as well as overlap between APOBEC3B and 3C, APOBEC3C and 3G, and APOBEC3B and 3G (**Fig. 2c; Extended Data Fig. 2d)**. Subsequent gene set enrichment analyses of significantly differentially expressed genes (gProfiler) revealed disruption of multiple pathways involved in RNA metabolism and RNA splicing regulation in APOBEC3B, APOBEC3C, and APOBEC3G overexpressing HSPCs compared to pCDH backbone-transduced controls (**Fig. 2d; Supplementary Table 2**). Notably, changes in these pathways are characterized by overexpression of genes encoding splicing factors, including U2AF1, ZRSR2 and RBM17, several SRSF family members (SRSF1, SRSF2, SRSF9), spliceosome component SF3B4, and multiple HNRNPs, all known to be crucial for hematopoiesis and implicated in CH (**Fig. 2e**).^38,41–44^ We also detected differential splicing, characterized by exon skipping, across all APOBEC3 family member overexpression conditions compared to backbone pCDH transduced HSPCs (**Fig. 2f; Extended Data Fig. 2e-g; Supplementary Table 3**). Specifically, in APOBEC3B, APOBEC3C, and APOBEC3G overexpressing cells, we detected 408, 359, and 424 exon skipping events involving 368, 310, and 355 genes, respectively. For these analyses, we utilized a false discovery rate (FDR) threshold of <0.05 and a percent spliced-in (PSI) value of >0.1. Interestingly, each APOBEC3 family member predominantly affected a distinct set of genes (**Fig. 2e; Extended Data Fig. 2f-g**).

RNA splicing is regulated by both trans- and cis-regulatory elements. The trans-regulatory elements include RNA binding proteins, which were found to be differentially expressed upon APOBEC3B, APOBEC3C, and APOBEC3G lentiviral transduction (**Fig. 2e**). The cis-regulatory elements comprise donor and acceptor splice sites, the branchpoint sequence, and various RNA-binding motifs, such as enhancers and silencers. However, our combined HSPC WGS and RNA-seq analyses revealed no significant differential APOBEC3-mediated C-to-U RNA editing surrounding the intron-exon junctions where both splice sites and branchpoint sequences are located (**Extended Data Fig. 2h**). Similarly, we did not detect enrichment of APOBEC3-mediated SBS near splice junctions (**Extended Data Fig. 2i**). Thus, distinct splicing patterns observed upon APOBEC3 lentiviral transduction may be primarily driven by differential expression of genes important for RNA splicing regulation, i.e., the trans-regulatory elements.

### APOBEC3C induces exon skipping in aged bone marrow HSPCs

Because exon skipping has been linked to leukemia stem cell generation^31,42,45,46^, we sought to further investigate the functional consequences of APOBEC3C activation in HSPCs. Among APOBEC3 family members examined, APOBEC3C uniquely induces both an increased mutational burden and widespread transcriptional changes, including alterations in RNA splicing. Moreover, we previously observed HPC expansion in CB following lentivirally enforced APOBEC3C expression, further highlighting its potential to affect HSPC proliferation and drive CH.^35,47^ To determine whether these effects extend beyond CB HSPCs, we transduced CD34^+^ ABM HSPCs with APOBEC3C followed by WGS, RNA-seq, and functional analyses (**Fig. 3a; Extended Data Fig. 3a,b**).

**Figure 3:**
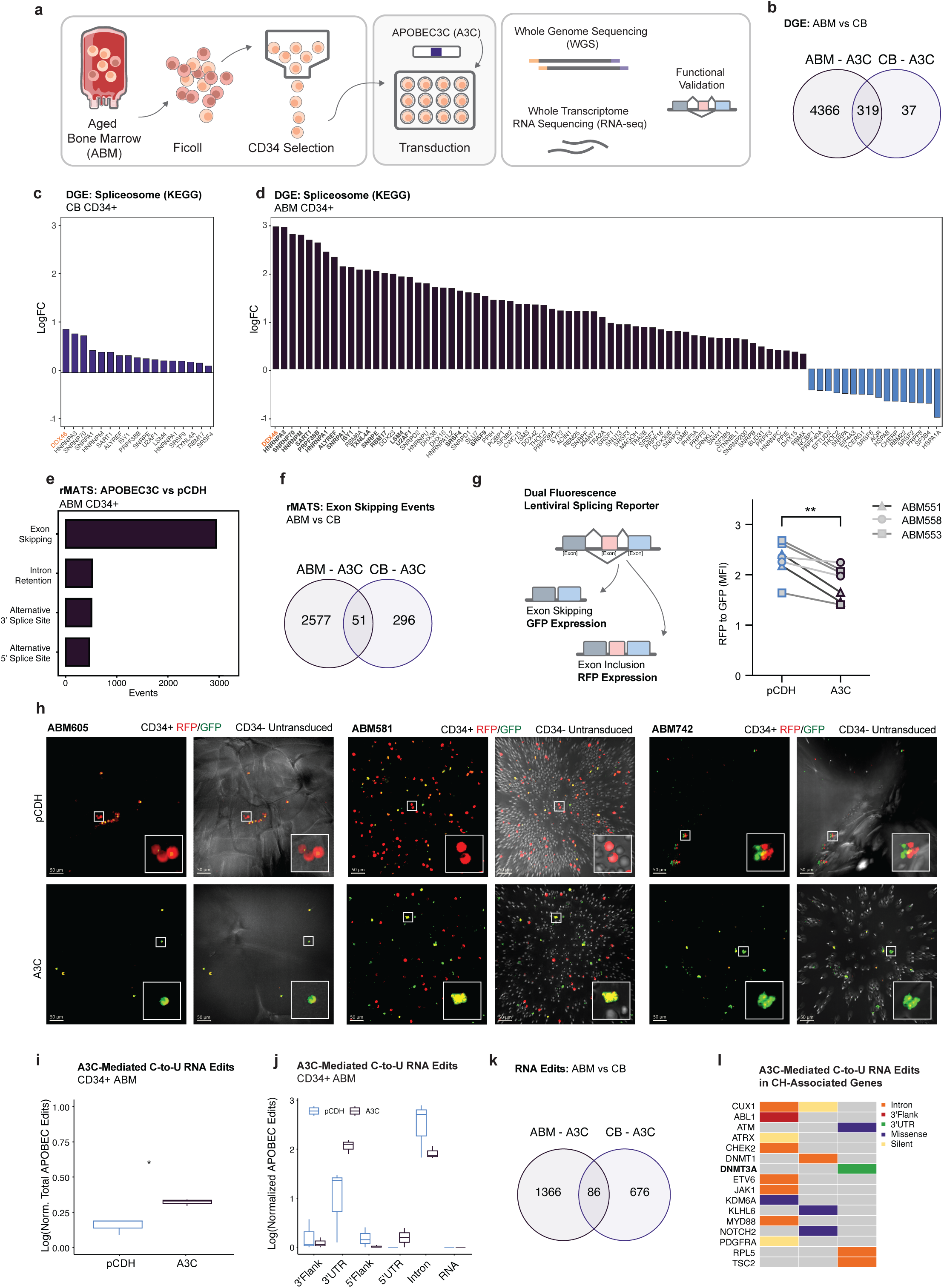
APOBEC3C induces exon skipping in aged bone marrow HSPCs **a.** Schematic representation of the experimental design in aged bone marrow (ABM) hematopoietic stem and progenitor cells (HSPCs). **b.** Venn diagram visualizing the number of significantly differentially expressed genes in ABM CD34⁺ cells transduced with APOBEC3C versus pCDH, and in cord blood (CB)-derived CD34⁺ cells transduced with APOBEC3C versus pCDH. **c.** Bar plot showing the fold changes (LogFC) of all significantly differentially expressed genes belonging to the KEGG Spliceosome pathway following APOBEC3C lentiviral transduction compared with pCDH in CB CD34⁺ HSPCs. **d.** Bar plot showing the fold changes (LogFC) of all significantly differentially expressed genes belonging to the KEGG Spliceosome pathway following APOBEC3C lentiviral transduction compared with pCDH in ABM CD34⁺ HSPCs. **e.** Quantification of differential splicing events, including exon skipping, intron retention, alternative 3’ splice site usage, and alternative 5’ splice site usage in ABM HSPCs cells lentivirally transduced with APOBEC3C compared to pCDH backbone transduced controls, using multivariate analysis of transcript splicing (rMATS). **f.** Venn diagram visualizing the number of exon skipping events in ABM CD34⁺ cells transduced with APOBEC3C versus pCDH, and in CB CD34⁺ cells transduced with APOBEC3C versus pCDH. **g.** Left: Schematic of lentiviral dual-fluorescence MAPT splicing reporter. Right: FACS analysis of GFP and RFP mean fluorescence intensity (MFI) in the live cell population of CD34^+^ selected cells derived from ABM following sequential lentiviral transduction with a splicing reporter and either APOBEC3C or control pCDH backbone. Error bars represent SD. Significance was determined using a paired t-test (p=0.0030) **h.** Representative fluorescence microscopy images of lentiviral dual-fluorescence MAPT splicing reporter transduced CD34⁺ ABM HSPCs, co-cultured with autologous untransduced CD34^-^cells and a porcine gelatin SURGIFOAM® matrix, which served as a physical scaffold. CD34^+^ HSPCs were sequentially lentivirally transduced with the dual-fluorescence MAPT splicing reporter and either APOBEC3C or the pCDH backbone control. **i.** Boxplot of total transcriptome-wide APOBEC3-mediated C-to-U RNA edits in ABM HSPCs following lentiviral transduction with pCDH backbone or APOBEC3C. **j.** Boxplot of transcriptome-wide APOBEC3-mediated C-to-U RNA edits in ABM HSPCs following lentiviral transduction with pCDH backbone or APOBEC3C per gene region; untranslated region (UTR). **k.** Venn diagram visualizing the number of C-to-U RNA edits in ABM CD34⁺ cells transduced with APOBEC3C versus pCDH, and in cord blood (CB) CD34⁺ cells transduced with APOBEC3C versus pCDH. **l.** Oncoplot illustrating APOBEC3-mediated C-to-U RNA edits in clonal hematopoiesis (CH)-associated genes following APOBEC3 lentiviral transduction of ABM HSPCs.

Differential gene expression detected 4685 genes that were significantly differentially expressed in APOBEC3C lentivirally transduced versus control HSPCs (**Fig. 3b**; **Supplementary Table 4;** adjusted p-value <0.05). Compared to CB HSPCs, we observed a 13.2-fold increase in differentially expressed genes in ABM HSPCs (**Fig. 3b**). In gene set enrichment analysis of significantly differentially expressed genes (gProfiler), the KEGG spliceosome pathway and Reactome RNA metabolism pathway emerged as top-ranked pathways (**Extended Data Fig. 3c**). We identified significant differential expression of several genes encoding RNA binding proteins including U2AF1, ZRSR2, MBNL1, PTBP1, RBFOX2, several SRSF family members (SRSF1, SRSF2, SRSF3, SRSF4, SRSF7, SRSF9), splicing factor RBM17, spliceosome components (SF3B2, SF3B4, SF3B5), and multiple HNRNPs, all known to be crucial for hematopoiesis and implicated in leukemia stem cell generation (**Fig. 3c,d; Extended Data Fig. 3d,e)** ^31,38,42^ We did not detect significant enrichment for both C-to-T DNA mutations and C-to-U RNA edits surrounding the intron-exon junctions in ABM (**Extended Data Fig. 3f**), further emphasizing the significance of APOBEC3C-mediated differential expression of genes encoding RNA binding proteins. Differential splicing analyses identified 4426 splicing events involving 2620 genes (**Fig. 3e**). APOBEC3C mainly induced exon skipping in ABM HSPCs compared to pCDH control HSPCs (2620/4426 events; **Fig. 3e,f**), with a 7.6-fold increase in exon skipping events compared to CB HSPCs (**Fig. 3f**), underscoring pronounced splicing dysregulation during aging.

To further investigate the role of lentiviral APOBEC3C expression on RNA splicing in ABM HSPCs, we examined the binding motifs of top differentially expressed RNA binding proteins. Interestingly, U2AF1, a component of the spliceosome, binds U2AF2, which recognizes and binds the polypyrimidine tract, representing a region rich in pyrimidine bases (cytosine and uracil) located upstream of the 3’ splice site. Following lentiviral APOBEC3C transduction, we observed enrichment of U2AF2 binding motif in intronic sequences (**Extended Data Fig. 3g**), suggesting that APOBEC3C-mediated base deamination introduces new binding sites. Accordingly, the mechanism underlying APOBEC3C-mediated regulation of splicing in ABM HSPCs might involve cytosine deamination, resulting in uracil incorporation into RNA. This process creates novel polypyrimidine tracts and could introduce new binding sites for RNA bindings proteins, which may have significant implications for splicing regulation.

Next, we utilized our lentiviral dual fluorescence splicing activity reporter capable of detecting exon skipping (GFP) and exon inclusion (RFP) in real-time at the single cell level (**Fig. 3g**).^48^ Using this dual fluorescence splicing reporter assay, we identified a significant decrease in the ratio of RFP to GFP, indicative of increased exon skipping (**Fig. 3g**). Confocal fluorescence microscopy analyses of CD34^+^ cells lentivirally transduced with the splicing reporter co-cultured with autologous untransduced CD34^-^cells and a porcine gelatin SURGIFOAM® matrix, which served as a physical scaffold in our stem cell niche nanobioreactor system^37^, similarly identified increased GFP signal upon lentiviral transduction with APOBEC3C, indicative of exon skipping, as well as interactions in the nanobioreactor niche between CD34^+^ cells transduced with the dual fluorescence splicing reporter and untransduced CD34^-^cells (**Fig. 3h, Video 1-2**). Together, these analyses support a previously unrecognized role for APOBEC3C-mediated splicing deregulation in human ABM HSPCs within the bone marrow niche.^48^

### APOBEC3C induces C-to-T mutations and RNA editing of CH-associated genes

To further investigate the contribution of APOBEC3C in CH, we performed HSPC WGS analyses that revealed increased SBS following lentiviral transduction of CD34^+^ ABM HSPCs with APOBEC3C versus pCDH controls **(Extended Data Fig. 3h-j**). In addition to pre-existing CH mutations, we identified likely pathogenic splice-site, multi-hit, frameshift, or missense mutations in *SMC1A, NOTCH1, ATM, ATRX, SMARCA4, NSD2, LUC7L2,* and *BCL11B* following APOBEC3C transduction of ABM HSPCs using single-sample mode (**Extended Data Fig. 3j)**. Subsequently, whole transcriptome C-to-U RNA editing analyses detected significantly increased C-to-U RNA edits in APOBEC3C overexpressing ABM HSPCs, compared with pCDH transduced controls, with enrichment of C-to-U editing at the 3’UTR (**Fig. 3i-k**). Targeted analyses of CH-associated transcripts revealed C-to-U RNA editing events in CUX1 in two out of three replicates; 3’UTR edits in DNMT3A; and missense edits in ATM, KLHL6 and NOTCH2 (**Fig. 3l**). Because CUX1 deregulation has been implicated in myeloid malignancy progression^49^, further studies will focus on the functional impact of CH-associated epitranscriptomic remodeling by APOBEC3C.

### APOBEC3C promotes progenitor replating and myeloid lineage skewing

Having characterized the genomic and epigenomic alterations associated with APOBEC3C overexpression, we investigated its broader functional impact on ABM HSPCs (**Fig. 4a**). In hematopoietic colony replating assays, APOBEC3C lentiviral transduction aberrantly increased replating capacity (**Fig. 4b,c; Extended Data Fig. 4a,b**). Consistent with increased regenerative capacity, lentiviral APOBEC3C-expressing HSPC colonies were larger than pCDH controls and predominantly consisted of myeloid colonies (**Extended Data Fig. 4a,b**).

**Figure 4:**
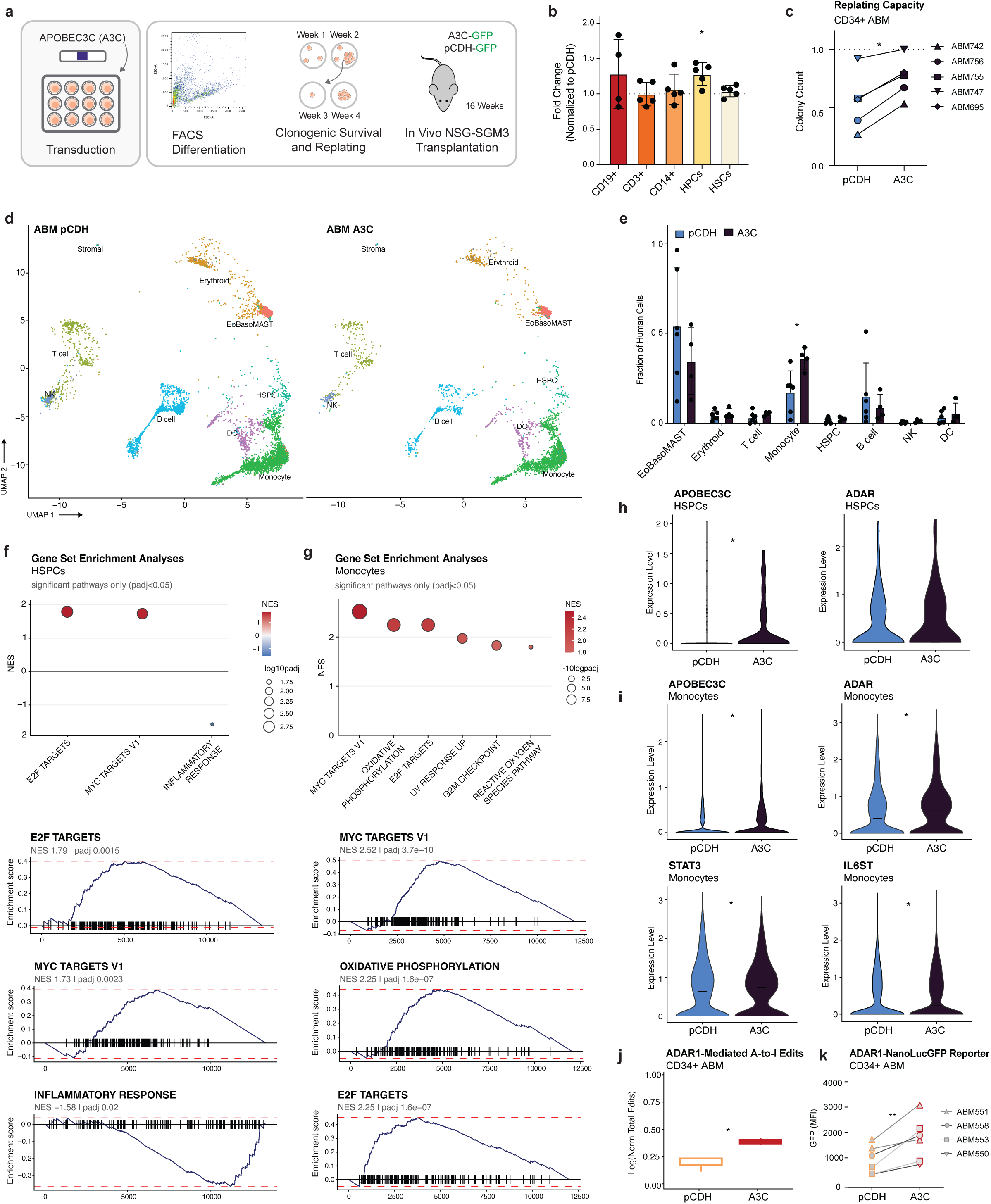
APOBEC3C promotes hematopoietic progenitor proliferation and myeloid skewing **a.** Schematic representation of the experimental design in aged bone marrow (ABM) hematopoietic stem and progenitor cells (HSPCs). **b.** ABM CD34^+^ cells were lentivirally transduced with APOBEC3C or a lentiviral backbone control. After 72h, cell counts were determined using FACS analysis. Error bars represent SD. Statistical significance was determined using a one-sample Wilcoxon signed-rank test against the hypothesized value of 1 (p=0.0160) **c.** Replating capacity was assessed and expressed as the percentage of replated colonies that grew out relative to the number plated. Error bars represent SD. Statistical significance was determined using a paired t-test (p=0.0263). **d.** UMAP visualization of single cell RNA sequencing (scRNA-seq) data of human bone marrow nuclear cells harvested upon engraftment of ABM HSPCs lentivirally transduced with pCDH (left) or APOBEC3C (right) **e.** Frequencies of predicted cell types based on the BoneMarrowMap reference atlas of human bone marrow hematopoiesis. Only mice with >1% human cells detected by scRNA-seq were included in the analysis. Error bars represent SD. Statistical significance was determined using a Mann–Whitney U test with individual mice as biological replicates (p=0.019). **f.** Top: Gene set enrichment analysis (GSEA) of HSPCs using the Molecular Signatures Database (MSigDB). Only mice with >1% human cells detected by scRNA-seq were included in the analysis. All pathways with a Benjamini–Hochberg-adjusted P value <0.05 are shown. Bottom: GSEA enrichment plots for the top three pathways enriched following lentiviral transduction with APOBEC3C compared with pCDH control. **g.** Top: Gene set enrichment analysis (GSEA) of monocytes using the Molecular Signatures Database (MSigDB). Only mice with >1% human cells detected by scRNA-seq were included in the analysis. All pathways with a Benjamini–Hochberg-adjusted P value <0.05 are shown. Bottom: GSEA enrichment plots for the top three pathways enriched following lentiviral transduction with APOBEC3C compared with pCDH control. **h.** Violin plots of APOBEC3C (p=0.038) and ADAR1 (not significant) gene expression in HSPCs upon engraftment of ABM HSPCs lentivirally transduced with pCDH or APOBEC3C. Statistical significance was determined using a Wilcoxon rank-sum test. **i.** Violin plots of APOBEC3C (p=1.92e-13), ADAR1 (p=6.99e-15), STAT3 (1.22e-7) and IL6ST (p=0.001) gene expression in HSPCs upon engraftment of ABM HSPCs lentivirally transduced with pCDH or APOBEC3C. Statistical significance was determined using a Wilcoxon rank-sum test. **j.** Boxplot of ADAR1-mediated A-to-I RNA edits in ABM HSPCs following lentiviral transduction with pCDH backbone or APOBEC3C. **k.** FACS analysis of GFP mean fluorescence intensity (MFI) in the live cell population of CD34^+^ selected cells derived from ABM following sequential lentiviral transduction with our ADAR1-NanolucGFP reporter, which detects A-to-I editing, and either APOBEC3C or the control pCDH backbone. Error bars represent SD. Significance was determined using a paired t-test (p=0.0099).

Subsequently, we compared engraftment of ABM CD34^+^ HSPCs lentivirally transduced with APOBEC3C and pCDH backbone intravenously in sublethally irradiated inflammatory niche NSG-SGM mice. Upon 16 weeks of engraftment, bone marrow mononuclear cells were harvested, selected for human CD45, and sent for scRNA-seq analyses (**Fig. 4d; Extended Data Fig. 4c**). To characterize human hematopoietic engraftment, we developed a bioinformatic pipeline to detect hematopoietic cell populations and track GFP-positive lentivirally transduced cells at single cell resolution. In engrafted mice, APOBEC3C lentiviral transduction induced a significant increase in monocytes and a trend toward decreased B cell frequencies compared to pCDH controls (**Fig. 4e**). Furthermore, GFP-based tracking demonstrated increased numbers of HSPCs and monocytes, alongside reduced B cell numbers indicative of myeloid skewing (**Extended Data Fig. 4d**). Although the frequencies of HSPCs were not significantly changed, differential gene expression analyses revealed enrichment of E2F and MYC target gene expression (**Fig. 4f**), consistent with increased regenerative capacity observed (**Fig. 4b**), alongside a decrease in inflammatory response-associated gene expression. In monocytes, we similarly observed increased expression of genes related to proliferation, as well as genes associated with stress responses (**Fig. 4g**). Furthermore, we confirmed significantly increased APOBEC3C expression in both HSPCs and monocytes and observed increased expression of genes involved in inflammatory signaling, including STAT3 and IL6ST, as well as ADAR1, potentially reflecting activation of cellular stress response pathways (**Fig. 4h,i**).

Then we determined ADAR1 expression levels in our whole transcriptome ABM and CB HSPC datasets. Interestingly, APOBEC3C lentiviral transduction significantly decreased the ADAR1 isoform ratio, specifically reducing ADAR1p150 expression, in CB HSPCs (**Extended Data Fig. 4e-g**). In ABM HSPCs, significantly increased levels of ADAR1-mediated A-to-I editing were detected by whole transcriptome A-to-I RNA editing analyses (**Fig. 4h-j**). This was confirmed by intracellular ADAR1p150 protein FACS analysis (**Extended Data Fig. 4j**), along with ADAR1-NanoLuc-GFP activity reporter assays, which showed elevated GFP levels indicative of increased A-to-I editing in ABM HSPCs (**Fig. 4k**).

### Increased C-to-U RNA editing in myeloproliferative neoplasm hematopoietic stem cells

Our previous studies revealed increased expression of APOBEC3C and ADAR1p150 in MPN HSCs and HPCs, stress-induced APOBEC3C activation and ADAR1p150 self-renewal gene expression upon inflammation, microgravity, and fight-or-flight responses, ADAR1-mediated A- to-I RNA editing signatures, and mutational profiles that conformed with an APOBEC3C-driven mutagenesis signature in HSPCs.^35–37,50^ However, APOBEC3-mediated C-to-U RNA editing in MPNs has not been investigated yet. Using our bioinformatic pipeline to detect C-to-U editing using whole transcriptome sequencing analyses, we discovered increased APOBEC3-mediated RNA edits specifically in HSCs among patients diagnosed with polycythemia vera (PV), essential thrombocythemia (ET), and myelofibrosis (MF; **Fig. 5a; Extended Data Fig. 5a**). Notably, editing events occurred in similar genomic regions across different patients, with editing patterns clearly distinct from those observed in ABM, particularly in HPCs (**Extended Data Fig.5a**). Among APOBEC3 family members, APOBEC3C expression showed the strongest positive association with total C-to-U RNA edits (**Extended Data Fig. 5b**). Notably, the top 50 edited genes included CUX1, a CH-associated gene. Targeted analysis of C-to-U editing in CH-associated genes further identified editing events that distinguished MPNs from ABM, including missense and multi-hit edits in CUX1 and TSC2 in a substantial proportion of patients (**Fig. 5b**). Missense edits in CUX1 predominantly occurred in exon 11 (**Fig. 5c**). Although their functional and clinical significance remains unknown and warrants further investigation, these edits may represent potential biomarkers. Moreover, AML patients with high APOBEC3C expression levels showed decreased overall survival (**Extended Data Fig. 5c**), further supporting its potential as a biomarker.

**Figure 5:**
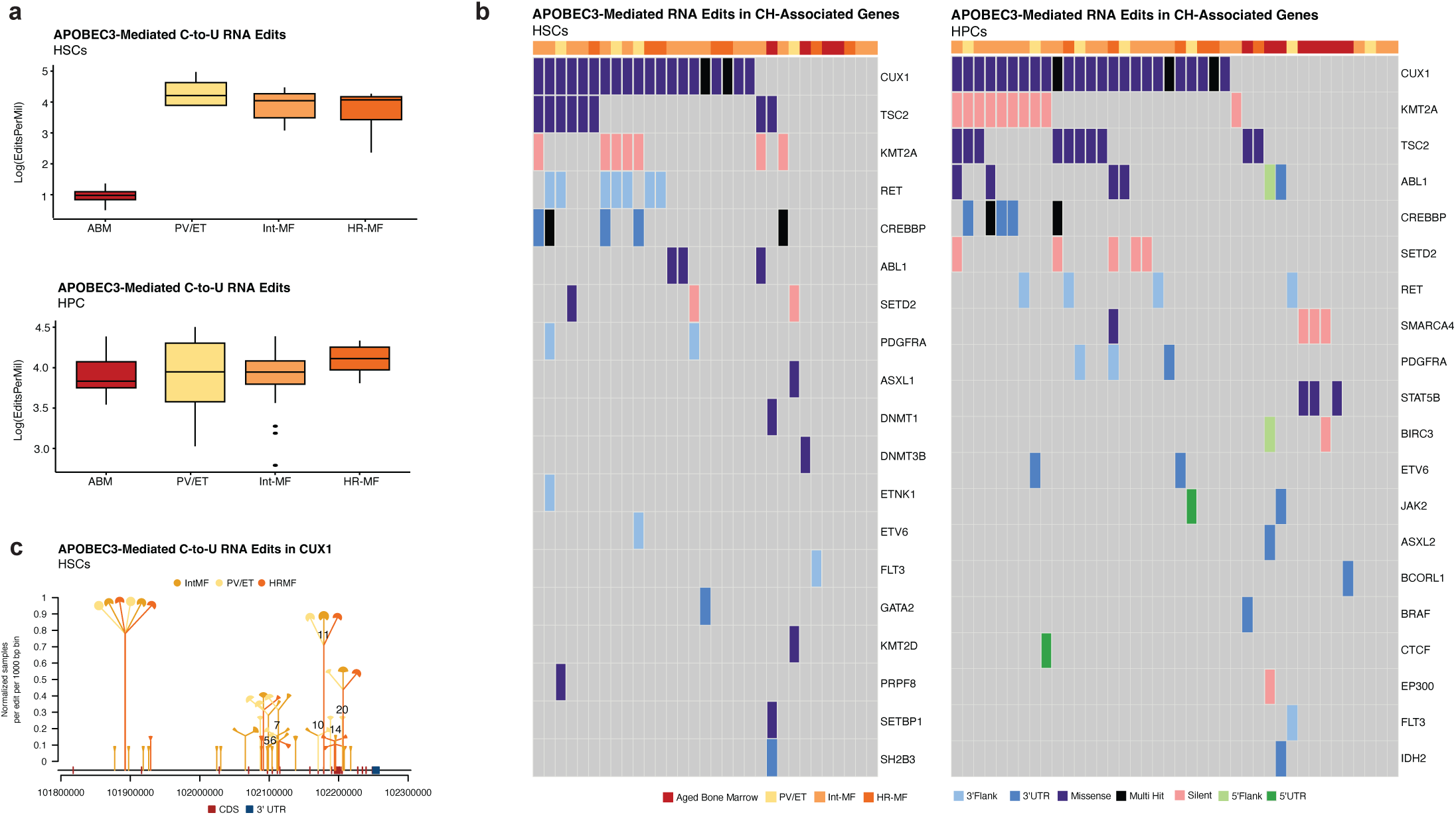
Increased C-to-U RNA editing in myeloproliferative neoplasm hematopoietic stem cells **a.** Boxplot of total transcriptome-wide C-to-U RNA edits in aged bone marrow (ABM), polycythemia vera (PV), essential thrombocythemia (ET), and National Comprehensive Cancer Network (NCCN) guideline-defined intermediate-risk (Int-MF) and high-risk (HR-MF) myelofibrosis. Data are shown for FACS-purified hematopoietic stem cells (HSCs; top) and hematopoietic progenitor cells (HPCs; bottom). **b.** Oncoplot illustrating C-to-U RNA edits in clonal hematopoiesis (CH)-associated genes in ABM, PV, ET, Int-MF and HR-MF for HSCs (left) and HPCs (right); untranslated region (UTR) **c.** Dandelion plot of C-to-U RNA edits in CUX1 across HSCs from patients diagnosed with PV, ET, Int-MF and HR-MF; coding sequence (CDS), untranslated region (UTR)

Additional investigation into the impact of APOBEC3C was performed using lentiviral shRNA-mediated knockdown. We first performed titration studies in the interferon-responsive TF1a leukemia cell line to validate dose-dependent APOBEC3C knockdown using our shRNA construct (**Extended Data Fig. 5d**). Notably, we also observed decreased STAT3 expression and reduced ADAR1 activation, as indicated by the ADAR1-NanoLucGFP reporter assay (**Extended Data Fig.5e-g**). These findings support a functional connection between APOBEC3C expression and STAT3/ADAR1-associated signaling in hematologic malignancies, while its potential as a therapeutic target needs to be determined in future studies.

## Discussion

Building on our recent findings implicating APOBEC3C in accelerated HSPC aging^36,37^ and MPN disease progression^35^, we investigated the functional hierarchy of APOBEC3 family members on DNA mutational burden, epitranscriptomic alterations, and hematopoietic function in HSPCs, and revealed that lentiviral transduction of APOBEC3C uniquely drives both genomic and epitranscriptomic changes in CB and ABM HSPCs. Most prominently, these changes included increased C-to-U RNA edits, increased exon skipping, and altered expression of RNA splicing factors previously implicated in CH,^38^ suggesting that APOBEC3C promotes widespread epitranscriptomic remodeling, ultimately enhancing replating capacity and promoting age-associated myeloid differentiation bias.

Genome-wide WGS analyses revealed that APOBEC3 family members differ in their capacity to induce DNA mutations. Compared to APOBEC3B, 3D, and 3G, APOBEC3C and 3F exhibited a higher mutational burden, showing close resemblance to MPN HSPCs.^35^ Mutational signature analysis identified SBS1 and SBS5, associated with C-to-T mutations predominantly at CpG sites and age-related mutational processes.^40^ Although SBS2 and SBS13 are established as APOBEC3-associated signatures, their relevance to APOBEC3 activity in primary human HSPCs remains unclear. Notably, APOBEC3 expression does not necessarily predict APOBEC3-associated mutational activity, which can occur episodically^51^, and limited DNA editing activity reported using engineered APOBEC3 reporter systems in HEK293T^52^, may not reflect its activity in human HSPCs. Thus, while mutations were predominantly C-to-T, APOBEC3C deamination targets and binding preferences in human HSPCs warrant future investigations^18^.

We further discovered APOBEC3B, 3C, and 3G-induced RNA splicing dysregulation, with APOBEC3C having particularly pronounced effects in aged HSPCs. RNA splicing is a tightly regulated process, and even without splicing factor mutations, dysregulation is recognized as a hallmark of cancer^53,54^. Shifts in the balance of splicing factors or recognition of its binding site can profoundly alter splicing outcomes, often resulting in increased exon skipping^48^, consistent with what we observed in ABM HSPCs following lentiviral APOBEC3C transduction. While APOBEC3C-mediated RNA splicing deregulation was observed in both CB- and ABM HSPCs, its impact in ABM was approximately ten-fold greater, suggesting that ABM cells may harbor intrinsic vulnerabilities, making them more susceptible to APOBEC3C activation.

Beyond its effects on RNA splicing, APOBEC3C overexpression significantly increased C-to-U RNA editing in aged HSPCs, particularly in 3′UTRs, which can influence transcript stability and post-transcriptional gene regulation, revealing an additional layer of APOBEC3-mediated epitranscriptomic remodeling. These findings are particularly interesting in the context of previous work from our group demonstrating that ADAR1-mediated A-to-I editing regulates micro-RNA biogenesis, influencing malignant progenitor propagation.^55–57^ Furthermore, previous studies have established stress-induced APOBEC3 activation, with APOBEC3A driving widespread C-to-U editing in monocytes and macrophages under inflammatory and hypoxic conditions^58^, APOBEC3G mediating RNA editing in natural killer cells upon mitochondrial hypoxic stress^59^, and APOBEC3C activation in human HSPCs following inflammation, microgravity and ‘fight or flight’ responses^35–37^. Here, APOBEC3C overexpression induced stress responses in monocytes, alongside enhanced HSPC regenerative capacity. This potential feed-forward loop between stress-induced APOBEC3C activation and stress-associated hematopoietic remodeling may be particularly relevant to MPN, where stressors including chronic inflammation contribute to disease progression. In this context, TRIM22, an interferon-inducible E3 ubiquitin ligase involved in antiviral defense, was consistently downregulated following APOBEC3 overexpression and in MPN HPCs, and reflect broader alterations in innate immune-associated pathways, which seem of particular interest given reported retrotransposon activation in leukemia stem cells and MPNs.^60,61^ Consistent with a role in CH and precancer initiation, APOBEC3 expression was observed in chronic lymphocytic leukemia^62^, while APOBEC3-mediated mutagenesis has been implicated in CH and MDS following retroviral gene therapy.^63^

Lastly, we observed increased APOBEC3-mediated C-to-U RNA edits in MPN HSCs, which were previously found to have increased APOBEC3C expression.^35^ Consistent with this observation, APOBEC3C expression showed the strongest positive association with RNA editing burden among the APOBEC3 family members examined. Notably, C-to-U RNA edits distinguished MPN from ABM and included genes implicated in CH. Loss-of-function mutations in CUX1 have been associated with clonal expansion of HSPCs and increased risk of AML transformation.^49^ We observed C-to-U RNA edits in CUX1 following lentiviral APOBEC3C overexpression in 2/3 CB and 2/3 ABM HSPC replicates, 25/32 MPNs, whereas these events were not detected in primary ABM HSCs or HPCs. Though, the functional and clinical significance of these edits require future studies. Thus, our findings provide new insight into how APOBEC3C-mediated epitranscriptomic remodeling may contribute to age-associated CH and precancer initiation, support its potential as a biomarker, and provide a foundation for exploring APOBEC3C as a precancer interception therapeutic target.

## Methods

### Human Subjects

Aged bone marrow was obtained from consented individuals undergoing knee or hip replacement surgery at Scripps Health according to Institutional Review Board-approved protocol (# IRB-20-7692). Patient characteristics of specimens used in this study can be found in **Supplementary Table 6**. Peripheral blood and bone marrow samples from patients diagnosed with MPNs were obtained from consenting patients at the University of California, in accordance with a UC San Diego Human Research Protections Program Institutional Review Board-approved protocol (#131550). Human cord blood was purchased as purified CD34^+^ cells (AllCells).

### Cell culture

Human cell lines TF1a and HEL were cultured at 37 °C in RPMI supplemented with 10% FBS and maintained according to ATCC protocols. All cell lines were confirmed to be mycoplasma-free with repeated testing.

### Patient Sample Processing

Peripheral blood mononuclear cells were isolated by Ficoll-paque density centrifugation and cryopreserved in liquid nitrogen. CD34^+^ cells were selected from peripheral blood mononuclear cells from both MPN patients and normal controls by magnetic bead separation (MACS; Miltenyi, Bergisch Gladbach, Germany). Cells were used fresh for functional experiments.

### APOBEC3C lentiviral vectors and transduction

Lentiviral plasmid backbone (pCDH-EF1-MSC-T2A-coGFP) was purchased (System Biosciences, Palo Alto, California, UCA) and human APOBEC3B, C, D, F and G were cloned into this vector by Eton Bioscience (San Diego, California, USA). Additionally, we designed a pCDH-EF1-MSC-T2A-coCFP plasmid backbone with APOBEC3C. Lentivirus was produced by GT3 Salk Core (San Diego, California, USA). Lentivirus used for shAPOBEC3C and shControl transduction was obtained from Vectorbuilder (Chicago, Illinois, USA). All lentivirus was tested by transduction of HEK293T and TF1a cells and efficiency was assessed by FACS analysis of GFP or CFP signal, and qRT-PCR. Lentiviral transduction of primary cells was performed at a MOI of 100-200. The cells were cultured for 72 hours in 96-well plate (5e10^5^ cells per well) containing StemPro-34 SFM medium supplemented with a cytokine cocktail consisting of SCF (50ng/ml), TPO (10ng/ml), FLT3 (50ng/ml), and IL-6 (10ng/ml)^47,57,64,65^. The transduced cells were collected after 72 hours of incubation, either into RLT for RNA extraction, pelleted for DNA extraction, or processed directly for functional assays.

### Whole-genome sequencing

DNA from the peripheral blood CD34^+^ population was extracted according to manufacturer recommendations using QIAamp DNA Blood Mini Kit (QIAGEN, Cat# 51104). Extracted DNA from these experiments was sent to Novogene (Sacramento, California, USA) for whole genome sequencing. DNA sequencing library preparation was performed using the NEBNext® DNA Library Prep Kit (New England Biolabs) following the manufacturer’s recommendations. Qualified libraries were paired and sequenced on an Illumina HiSeq X (CB) or Illumina NovaSeq 6000 platform (ABM) using 150 PE sequencing chemistry to 90X coverage according to effective concentration and data volume.

### Identification of somatic mutations from whole-genome bulk sequencing

Raw sequence data were downloaded to the Triton Shared Computing Cluster^66^ from ftp server link shared by Novogene (Sacramento, CA). All the post-sequencing analyses were performed within TSCC at UC San Diego. The methodology for identification of somatic mutations from bulk sequencing data follows established approaches from large genomics consortia^19^. Briefly, quality assurance of the raw FASTQ files were evaluated using FastQC (Version 0.12.0) and Mosdepth (Version 0.3.4)^67,68^. Raw sequence reads were aligned to the human reference genome GRCh38.d1.vd1. The aligned reads were marked duplicated using MarkDuplicates (Picard) from GATK (Version 4.0.11.0)^69^. Concordance between paired samples were checked using Conpiar version 0.274. An ensemble variant calling pipeline (EVC) was used to identify single nucleotide variants (SNV) and short insertions and deletions (indels). EVC implements the SNV and indel variant calling from four variant callers (Mutect2, Strelka2, Varscan2, and MuSE2) and only mutations that are identified by any two variant callers were considered as bona fide mutations^69–72^.

### Analysis of mutational profile

Analysis of mutational profiles was performed using our previously established methodology with the SigProfiler suite of tools used for summarization, simulation, visualization, and assignment of mutational profiles. Briefly, mutational matrixes for SBS, DBS and Indels were generated with SigProfilerMatrixGenerator (Version 1.2.16)^73^.Plotting of each mutational profile was done with SigProfilerPlotting (Version 1.3.13). Germline variants were filtered against the dbsnp138 database with the criterion that the alternate Common Allele Frequency >= 0.02.

### Annotation of the mutations

The mskcc/vcf2maf (Cyriac Kandoth)^74^ was used to annotate each mutation with its genomic coordinates using Ensembl Variant Effect Predictor (VEP, v106) and the annotated files were saved as mutation annotation format (MAF). Plotting of the genetic annotation was done using Maftools^75^ (version 2.16.0) within R (v4.3.1) statistical language. Annotation of each C>T mutation to its CpG island is done using MutationalPatterns (v3.10.0) packages in R. Venn diagram of overlapping mutations was done using VennDiagram (v1.7.3) package in R.

### APOBEC3-associated analysis

For germline-filtered mutations, APOBEC-associated mutations were extracted based on whether they fall under the T[C>T/G]W context in SBS96. The trinucleotide context information was generated by Sigprofiler (v1.3.5) and was intersected with the germline-filtered vcf files using bedtools (v2.31.1). Contingency tables were generated by averaging three replicates for each sample. Statistical significance was evaluated using Fisher’s exact test on these contingency tables. For each of the all-mutation and APOBEC-associated-mutation categories, vcf2maf (v1.6.22) and ensembl-vep v106.0 were run to obtain annotated maf files. Finally, oncoplots for non-synonymous mutations were plotted for each sample using maftools v2.24.0 within R v4.5.0. Different variant allele frequencies and genes of interest were selected for each oncoplot. Variant classifications included: Frame_Shift_Del, Frame_Shift_Ins, Splice_Site, Translation_Start_Site, Nonsense_Mutation, Nonstop_Mutation, In_Frame_Ins, In_Frame_Del, Missense_Mutation, 3’Flank, 5’Flank, 3’UTR, 5’UTR, RNA, Silent, Intron, Splice_Region, and IGR.

### CH-associated analysis

To identify mutations affecting CH-associated genes in ABM HSPCs mutations were called using GATK Mutect2 single samples mode by using mark duplicated reads files from EVC pipeline, which is freely available under the permissive 2-clause BSD license at: https://github.com/AlexandrovLab/EnsembleVariantCallingPipeline. Variants were filtered using FilterMutectCalls, and only PASS variants were retained. These were annotated with Funcotator to generate MAF files. Variants within 123 genes associated with clonal hematopoiesis (CH) were selected for downstream visualization using Maftools (version 2.16.0) within R (v4.3.1) statistical language.” Additionally, identified variants were filtered with the “AlphaMissense” annotation for “likely pathogenic” mutations based on the pathogenicity score within the tool^76^, of which results were addressed in the manuscript text.

### Whole transcriptome (RNA-seq) Analyses

RNA-seq (2 x150bp) paired-end reads were generated and analyzed as described previously^48^. Briefly, following quality control with FastQC^77^ demultiplexed fastq files were aligned to the human genome, hg38, with STAR (v2.5.3a)^78^ with gene/transcript abundance quantified by RSEM (v1.3.0)^79^ GENCODE annotation (gencode.v43.annotation.gtf). Using functions from the R Bioconductor package edgeR^80^ and limma^81^ gene and isoform levels were separately aggregated for all samples, filtered for low expressing genes (counts per million < 1 in 50% of samples), Trimmed Mean of the M-values (TMM) normalized, and converted to log-transformed counts per million (lcpm) values for plotting. Enrichment analysis was performed with gProfiler (https://biit.cs.ut.ee/gprofiler/gost) using KEGG, Reactome, and WikiPathway ontologies and an adjusted p-value threshold of 0.05.

### Differential Splicing Analyses

Bam files generated by STAR alignment to hg38 (described above) were used as input into the rMATS software (v4.1.0)^82^ to determine significant differential splicing events. For the event types Alterative 3’ Splice (A3’SS), Alternative 5’ Splice Site (A5’SS), Retained Intron (RI), and Skipped Exon (SE), events with FDR < 0.5 and |PSI| > 0.1 using both junction counts and exon counts were considered significant.

### RNA Editing Analyses

RNA editing events within each sample were determined as described previously^45^. ADAR-mediated events were assessed for which positive strand A to G and negative strand T to C variants were retained for quantification of total edits within each sample group. APOBEC3-driven events were the focus of this analysis. Similar to the ADAR-mediated events, mutations from C to T on the + strand (G to A on the – strand), were retained and annotated for summary analyses.

### Single-Cell RNA-Sequencing Analyses

Single-cell libraries were prepared using 10X-3’ v4 Gene Expression (10X Genomics) kit according to manufacturer’s instructions. The libraries’ quality and quantity were assessed using the TapeStation 4200 DNA High Sensitivity analysis and the Qubit 4 dsDNA high sensitivity assay. Cells were pooled and sequenced on the Illumina NovaSeq X platform (Novogene Corporation) using 25B flow cells and 1% PhiX.

Sequenced reads from each GEX library generated were aligned to the combined human and mouse reference genome, downloaded from the 10X genomics website (10X Genomics, GRCh38-and-GRCm39-2024-A) with the copGFP sequence from the lentiviral vector added as an additional reference to detect transduced engrafted cells. Quality control, alignment, and quantification of reads were performed using the Cell Ranger v.9.0.0 (10X Genomics) ‘count’ pipeline, generating count matrices that filtered out empty droplets. GEX count matrices were loaded with Seurat v.5.6.0^83^ and filtered for cells with outlier UMI counts (low quality cells and doublets), high mitochondrial gene expression (due to cellular stress or loss of cytoplasmic RNA), and low number of sequenced genes. Individual datasets were bioinformatically cleaned for doublets using DoubletFinder v2.0.6^84^ and count matrices were merged into a single object. Molecular count data for each sample were individually log normalized and scaled in Seurat with the additional steps of regressing UMI counts, and gene counts during scaling to mitigate their effects on dimensionally reduced clustering. Highly variable genes were identified as the union of the top 3,000 genes from each sample after normalization and then used to calculate principal components using RunPCA. Normalized and scaled samples were integrated using the IntegrateLayers function using the “scVIIntegration” method in Seurat.^83^ Visualization and clustering of the integrated object was performed by calculating the PCA for dimensionality reduction visualization using the UMAP function in Seurat.^83^ (After identifying clusters via FindClusters in Seurat, clusters with high mouse gene expression (>50%) were removed and remaining cells were filtered for >90% human gene expression and <10% mouse gene expression.

The reads from each GEX library were realigned to the human reference genome GRCh38 alone, downloaded from the 10X genomics website (10X Genomics, GRCh38-2024-A), and annotated with GENCODE v44 (Ensembl release 110) using the same Cell Ranger pipeline. These GEX count matrices were loaded with Seurat v.5.6.0^83^, subset to the barcodes of the clean human cells defined above, and merged into a single object which was normalized, scaled, PCA calculated, integrated, visualized by UMAP, and clustered following the methods described above. Cells were then projected onto the BoneMarrowMap reference atlas of human bone marrow hematopoiesis (v0.1.0)^85^ to predict immune cell types and visualize them by UMAP. These predicted cell types were then collapsed into common immune cell types for downstream analysis.

Differentially expressed genes (DEGs) were calculated using the ‘‘MAST’’ algorithm^86^ which is tailored to scRNA-seq data DEG analysis using a model that parameterizes both stochastic dropout and characteristic bimodal expression distributions, for the FindMarkers function of Seurat. DEGs from FindMarkers were used to generate ranked gene lists ordered by adjusted p-value change for Gene Set Enrichment Analysis (GSEA) using the fgsea v1.32.2.^87^ Fast gene set enrichment analysis and gene signatures were pulled from the Molecular Signatures Database (MSigDB) using msigdbr v[25.1.1],^88,89^ restricted to the Hallmark (H) and Reactome (C2:CP:REACTOME) collections. Pathways with a Benjamini-Hochberg adjusted p < 0.05 were considered significantly enriched. Normalized enrichment scores and running enrichment plots were visualized in R (version 4.4.3) using ggplot2 and the plotEnrichment function in fgsea.

### FACS Analyses

To assess cell types upon APOBEC3C expression, cells were stained with fluorescence antibodies against human CD34, CD38, CD14, CD19 and CD3. (CD34-BV421 (1:50), Biolegend; CD38-PE-Cy7 (1:50), BD; CD14-APC-Cy7 (1:50), Biolegend; CD3-FITC (1:50), Biolegend; CD19-AF700 (1:50), Biolegend) and 7AAD (Biolegend). Subsequently, we ran samples on the BD Fortessa X-20. To assess ADARp150 protein levels, cells were stained with viability dye (Thermo Fisher Scientific, Live/Dead Fixable NearIR Dead Cell Stain Kit) and CD34 (Biolegend, BV421, 1:50) prior to fixation and permeabilization for intracellular staining of human ADAR1 (Abcam, ADAR1-APC, 1:100)

### Dual Fluorescence Lentiviral Splicing Reporter

For the evaluation of in vitro splicing activity using a two-color fluorescence splicing reporter system, CD34^+^ selected cells from ABM were lentivirally transduced with the splicing reporter for 8-24 hours, followed by lentiviral transduction with either pCDH or APOBEC3C virus. Following 72 hours, cells stained with 7AAD were analyzed by FACS. Additionally, upon incubation, cells were co-cultured with untransduced CD34^-^cells derived from the same individual, in a single well glass bottom 35mm plate (MatTek Life Sciences Cat# P35G-0.170-14C) with a porcine gelatin SURGIFOAM® matrix (J&J MedTech) to function as a physical scaffold (trabecular bone). Subsequently, cells were imaged using the Dragonfly confocal microscopy system (Andor Technology, Belfast, Northern Ireland) or Olympus FluoView FV10i for real-time single cell assessment of splicing activity. Dragonfly images were analyzed by the Imaris software by Oxford Instruments.

### ADAR1 Reporter Assay

For RNA editing activity assays, CD34^+^ selected cells from ABM as well as cell lines TF1a and HEL, were lentivirally transduced with ADAR1 NanoLuc-GFP reporter for 24 hours, followed by lentiviral transduction with either pCDH, APOBEC3C, shControl or shAPOBEC3C virus. Following 72 hours, cells stained with 7AAD were analyzed by FACS.

### Colony Formation Survival and Replating Assay

Colony forming unit (CFU) survival and replating assays were performed as previously described^35^. Briefly, following 72 hours of lentiviral transduction with either pCDH, APOBEC3C, shControl or shAOBEC3C virus, 1,000 CD34^+^ cells were collected, and plated in in triplicates in methylcellulose (MethoCult™ H4335 Cat# 04330, Stem Cell Technologies Inc.). After two weeks of culturing primary colonies were counted. Subsequently, a minimum of 10 individual colonies were plucked per replicate, resuspended, and replated in fresh methylcellulose. Secondary colonies derived from replated individual colonies were counted after another 14 days of culturing.

### Humanized Engraftment Studies

All mouse studies were completed in accordance with University Laboratory Animal Resources and Institutional Animal Care and Use Committee of the University of California regulations. Immunocompromised NSG-SGM3 mice were bred and maintained in the Sanford Consortium for Regenerative Medicine vivarium according to IACUC approved protocols. Upon lentiviral transduction, 90,000 cells were transplanted by intravenous injection into 5-week-old NSG-SGM3 mice that had been irradiated with 150cGy using a 137Cs irradiator (Shepherd & Associates, San Fernando, CA, USA). Upon 16 weeks of engraftment, mice were humanely euthanized, followed by collection of bone marrow, spleen, and peripheral blood. Human CD45^+^ cells were selected from bone marrow mononuclear cells by magnetic bead separation (MACS; Miltenyi, Bergisch Gladbach, Germany), followed by scRNA-seq analyses.

### Kaplan-Meier Plots

Kaplan-Meier analyses of overall survival (OS) of acute myeloid leukemia (AML) patients stratified by APOBEC3C gene expression were performed using GEPIA, an interactive web server containing RNA sequencing data from TCGA and GTEx databases^90^. The RNA-Seq datasets GEPIA used is based on the UCSC Xena project (http://xena.ucsc.edu), which are computed by a standard pipeline. GEPIA uses the Log-rank test, also known as the Mantel–Cox test, for hypothesis testing. A median expression threshold is set as the cutoff for high-expression and low-expression cohorts. Cutoff-high expression and cutoff-low expression are set to 50%. Samples with an expression level higher than 50% are considered the high-expression cohort. Samples with an expression level lower than 50% are considered the low-expression cohort. The Cox proportional hazard ratio and the 95% confidence interval (CI) information are included in the survival plot. 95% CI is added as dotted line. Analysis is performed using acute myeloid leukemia samples from the TCGA database, which includes 173 bone marrow-derived patient samples.

## Data availability

Whole-transcriptome RNA-seq, scRNA-seq and WGS data have been deposited at the database of Genotypes and Phenotypes (dbGaP) and are available via phs002228.v1.p1 and phs004195.v1.p1.

## Code availability

Somatic mutations in whole-genome sequencing data were identified using our ensemble variant calling pipeline, which is freely available under the permissive 2-clause BSD license at: https://github.com/AlexandrovLab/EnsembleVariantCallingPipeline. All other computational tools utilized in this publication have been mentioned in the methodology section and can be accessed through their respective publications.

## Author contributions

C.H.M.J, J.I, and I.v.d.W conceived the study. I.v.d.W., J.I., E.K., J.P., W.M., and C.E., designed and/or performed experiments and analyzed data. T.W. performed RNA-seq, RNA splicing, and RNA editing analyses and managed all transcriptomics data and deposition, supervised by I.v.d.W. and C.H.M.J. R.W. and S.N. performed all WGS and CH-associated mutation analyses and managed all genomic data and deposition, supervised by I.v.d.W, L.A., and C.H.M.J. C.H.M.J., A.K., I.v.d.W., and E.K., consented and collected all primary patient samples under an IRB-approved protocol. I.v.d.W and C.H.M.J. wrote the manuscript, which was reviewed and edited by all authors. C.H.M.J. supervised all aspects of the project.

## Additional information

Extended data Extended data Figures 1-5

## Supplementary information

Supplementary Table 1. Differential gene expression results of CB HSPCs lentivirally transduced with APOBEC3B, APOBEC3C, APOBEC3D, APOBEC3F or APOBEC3G compared to pCDH backbone, related to Figure 2.

Supplementary Table 2. gProfiler analysis of CB HSPCs lentivirally transduced with APOBEC3B, APOBEC3C or APOBEC3G compared to pCDH backbone, related to Figure 2.

Supplementary Table 3. rMATS derived exon skipping events of CB HSPCs lentivirally transduced with APOBEC3B, APOBEC3C, APOBEC3D, APOBEC3F or APOBEC3G compared to pCDH backbone, related to Figure 2.

Supplementary Table 4. Differential gene expression results of ABM HSPCs lentivirally transduced with APOBEC3C compared to pCDH backbone, related to Figure 3.

Supplementary Table 5. gProfiler analysis of ABM HSPCs lentivirally transduced with APOBEC3C compared to pCDH backbone, related to Figure 3.

Table S6. Patient characteristics

Video 1. Fluorescence microscopy of dual fluorescence splicing reporter transduced CD34^+^ ABM HSPCs, co-cultured with their untransduced CD34^-^cells and a porcine gelatin SURGIFOAM® matrix, which served as a physical scaffold.

---

**Extended Data Fig. 1:**
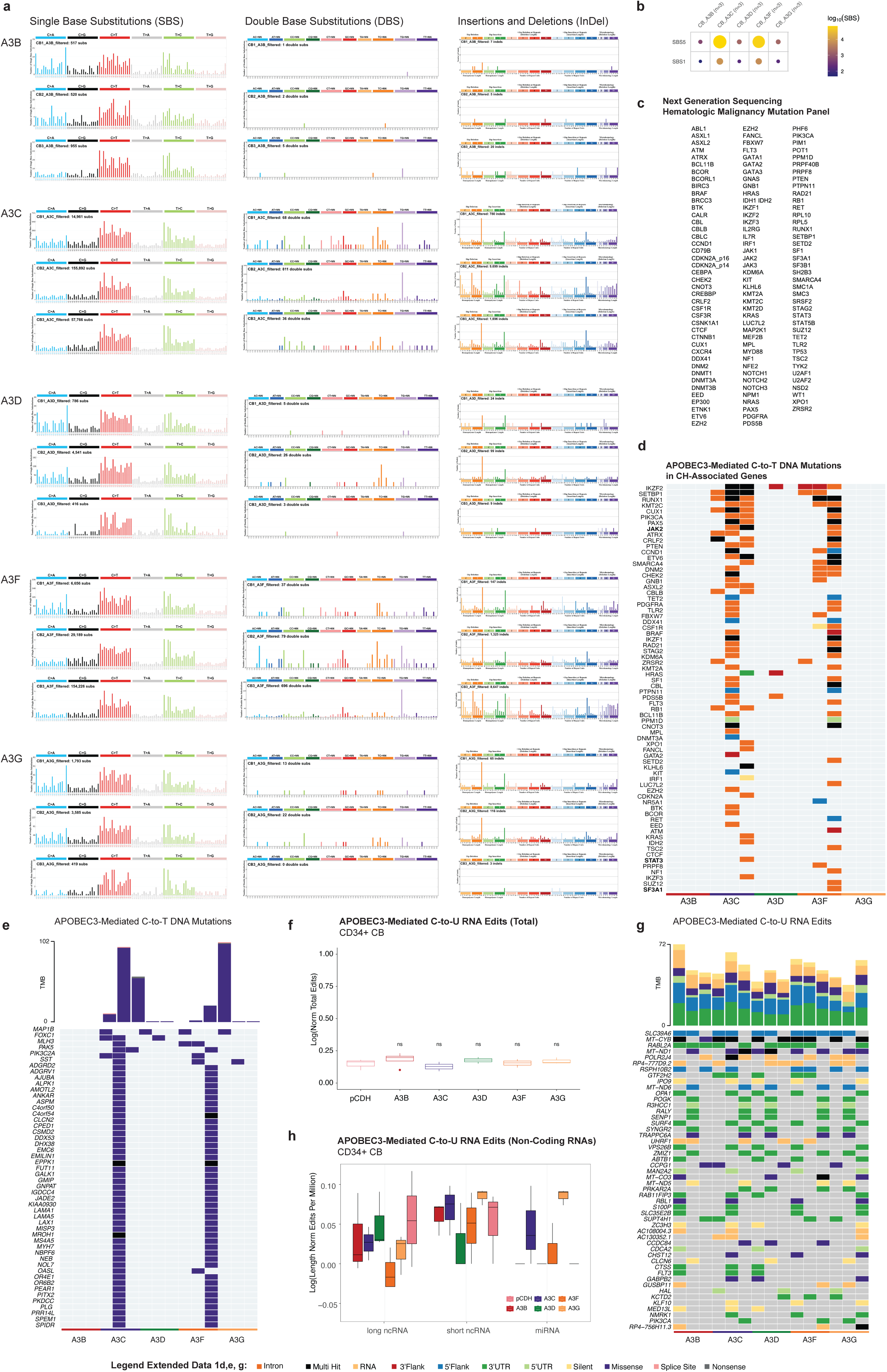
APOBEC3-induced epitranscriptomic and genomic editing in HSPCs. **a.** Mutational profiles across all three replicates of cord blood (CB) hematopoietic stem and progenitor cells (HSPCs) following lentiviral transduction with APOBEC3B, APOBEC3C, APOBEC3D, APOBEC3F, or APOBEC3G normalized to pCDH. Single base substitution (SBS) patterns are shown using the SBS96 classification (left), double base substitutions (DBS) using the DBS78 classification (middle), and small insertions and deletions (Indel) using the ID83 classification (right). **b.** Cosine similarity plotted for SBS COSMIC signatures SBS1 and SBS5 in APOBEC3-overexpressing CB HSPCs **c.** List of 125 clonal hematopoiesis (CH)-associated genes used for targeted mutational analyses. **d.** Oncoplot illustrating APOBEC3-mediated C-to-T DNA mutations in CH-associated genes following APOBEC3 lentiviral transduction of CB HSPCs. **e.** Oncoplot illustrating the top 50 genes affected by APOBEC3-mediated C-to-T DNA multi-hit or missense mutations following APOBEC3 lentiviral transduction of CB HSPCs. **f.** Boxplot of total transcriptome-wide APOBEC3-mediated C-to-U RNA edits in CB HSPCs following lentiviral transduction with pCDH backbone, APOBEC3B, APOBEC3C, APOBEC3D, APOBEC3F, or APOBEC3G. **g.** Oncoplot illustrating the top 50 genes affected by APOBEC3-mediated C-to-U RNA editing following APOBEC3 lentiviral transduction of CB HSPCs. **h.** Boxplot of APOBEC3-mediated C-to-U RNA edits in long and short non-coding RNAs (long ncRNAs), as well as micro-RNAs (miRNAs) in CB HSPCs following lentiviral transduction with pCDH backbone, APOBEC3B, APOBEC3C, APOBEC3D, APOBEC3F, or APOBEC3G.

**Extended Data Fig. 2:**
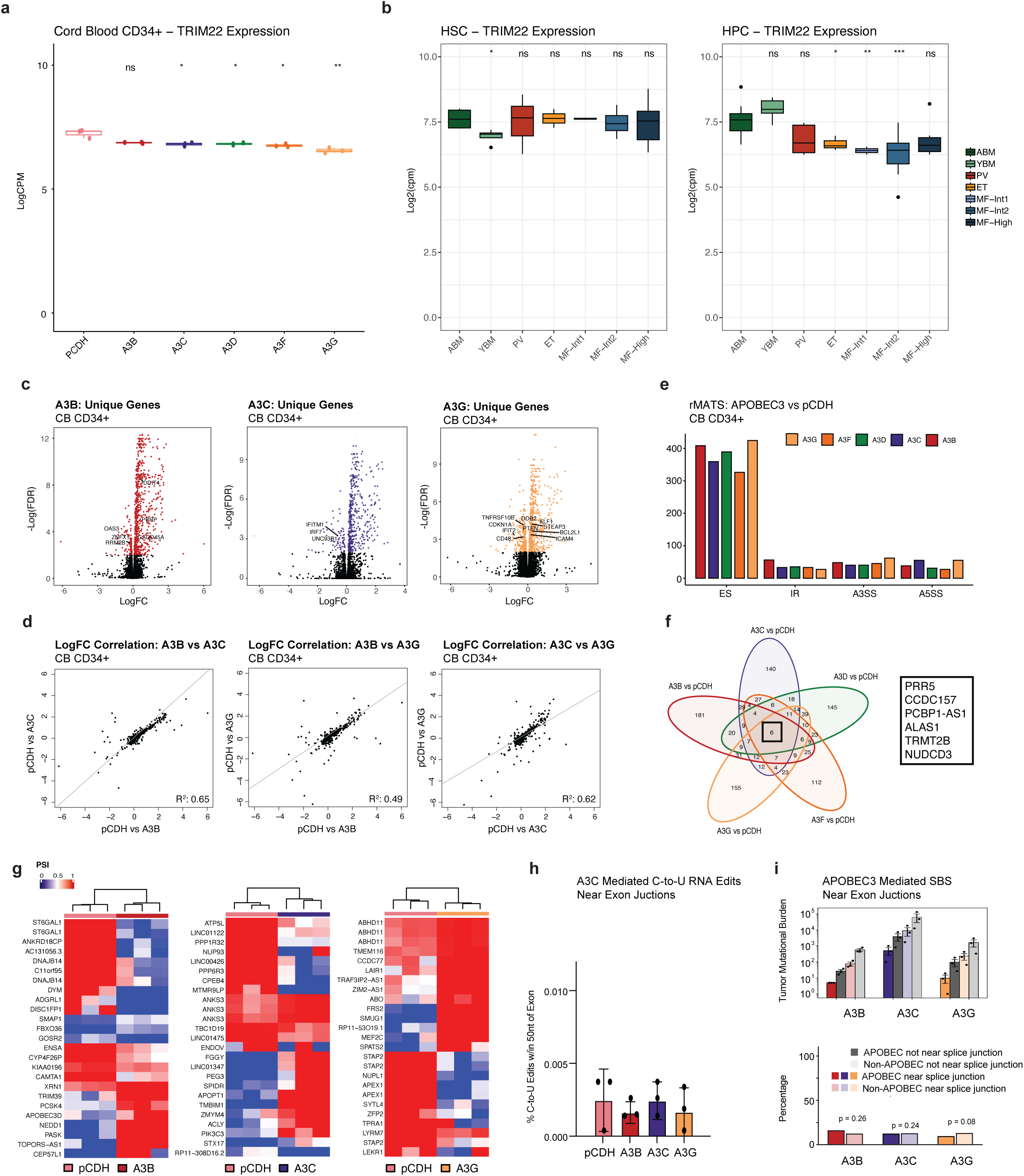
APOBEC3 activation decreases expression of the interferon-responsive TRIM22 gene and promotes splicing deregulation. **a.** RNA-seq based expression levels (log counts per million) of TRIM22 in cord blood (CB) hematopoietic stem and progenitor cells (HSPCs) lentivirally transduced with pCDH, APOBEC3B, APOBEC3C, APOBEC3D, APOBEC3F and APOBEC3G. **b.** RNA-seq-based expression levels (log counts per million) of TRIM22 in aged bone marrow (ABM), young bone marrow (YBM), polycythemia vera (PV), essential thrombocythemia (ET), and National Comprehensive Cancer Network (NCCN) guideline-defined intermediate-risk (Int-MF) and high-risk (HR-MF) myelofibrosis. Expression levels are shown for FACS-purified hematopoietic stem cells (HSCs; left) and hematopoietic progenitor cells (HPCs; right). **c.** Volcano plots of gene expression differences between APOBEC3B and pCDH, APOBEC3C and pCDH, and APOBEC3G and pCDH. Fold change (LogFC) is shown on the x axis with the negative log of the nominal p value on the y axis. Colored dots represent genes with nominal p<0.05. **d.** Correlation plots comparing fold changes (LogFC) of significantly differentially expressed genes following APOBEC3B versus APOBEC3C, APOBEC3B versus APOBEC3G and APOBEC3C versus APOBEC3G, each relative to pCDH control. **e.** Quantification of differential splicing events, including exon skipping (ES), intron retention (IR), alternative 3’ splice site usage (A3SS), and alternative 5’ splice site (A5SS) usage in CB HSPCs lentivirally transduced with APOBEC3B, APOBEC3C, APOBEC3D, APOBEC3F, or APOBEC3G, compared to pCDH backbone transduced controls, using multivariate analysis of transcript splicing (rMATS). **f.** Venn diagram visualizing the absolute number of exon skipping events induced by APOBEC3B, APOBEC3C, APOBEC3D, APOBEC3F, and APOBEC3G following lentiviral transduction in CB HSPCs compared to pCDH backbone transduced controls. Overlap of 6 genes exhibiting differential exon skipping events common to lentiviral transduction with APOBEC3B, APOBEC3C, APOBEC3D, APOBEC3F and APOBEC3G, include PR5, CCDC157, PCBP1-AS1, ALAS1, TRMT3B and NUDCD3. **g.** Heatmap showing the percent spliced in (PSI) values of the top 25 unique exon skipping events uniquely mediated by APOBEC3B, APOBEC3C, and APOBEC3G. Some genes are represented multiple times in the heatmap because they are affected by more than one exon skipping event. **h.** Boxplot of APOBEC3-mediated C-to-U RNA editing events in close proximity, within 50 nucleotides (nt), of exon-intron boundaries normalized to the total number of edits identified. No significant differences between conditions were observed. **i.** Absolute (top) and relative (bottom) numbers of APOBEC3-mediated single base substitutions (SBSs) within 50 nucleotides of intron-exon boundaries. Relative numbers are shown as SBSs near splice junctions and those not near splice junctions, normalized to the total number of SBSs in each category. No significant differences between conditions were observed.

**Extended Data Fig. 3:**
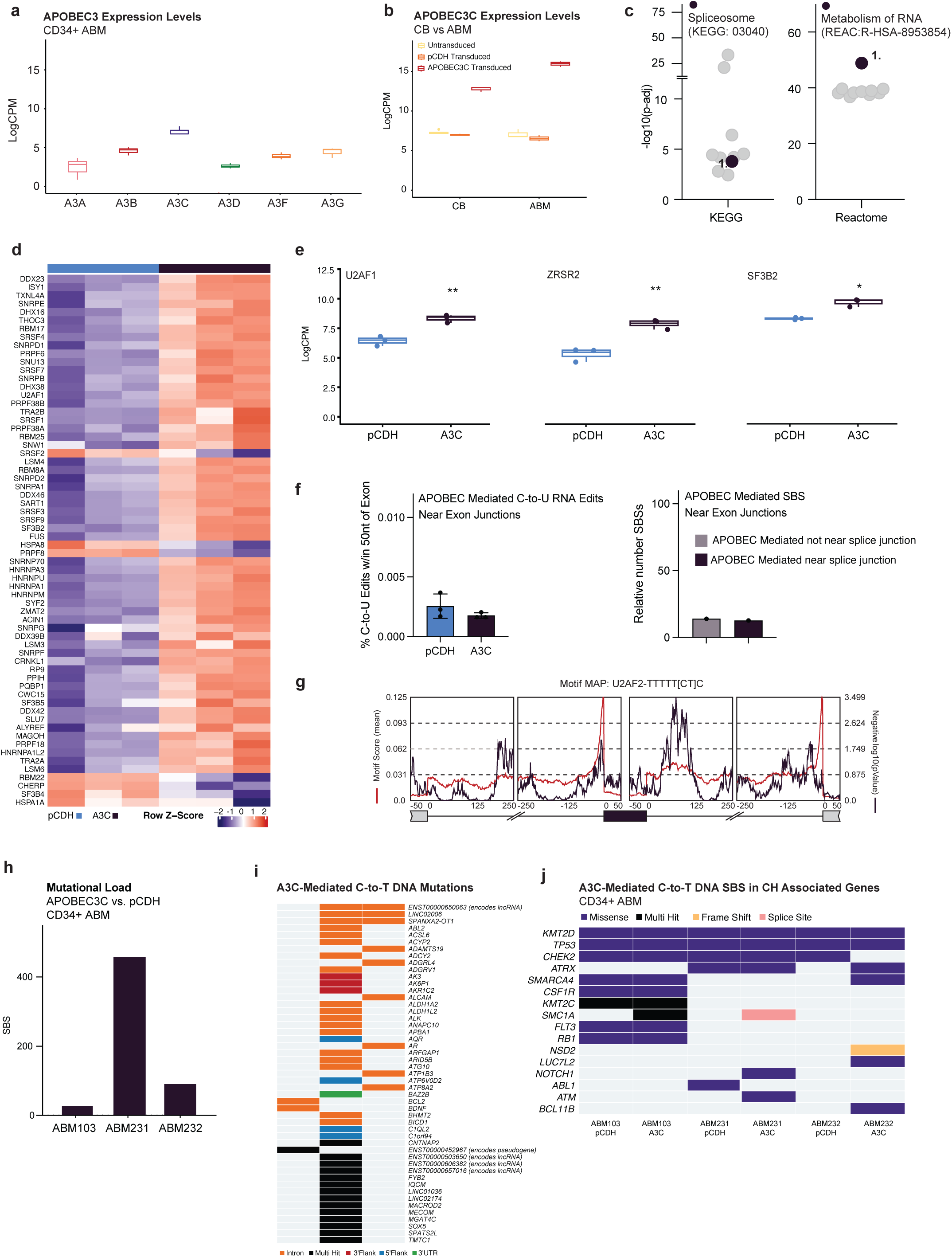
APOBEC3C induces differential expression of splicing factors and DNA mutations in aged bone marrow HSPCs. **a.** RNA-seq based expression levels (log counts per million) of APOBEC3A-H in aged bone marrow (ABM) hematopoietic stem and progenitor cells (HSPCs). **b.** RNA-seq based expression levels (log counts per million) of APOBEC3C in cord blood (CB) and ABM HSPCs upon lentiviral transduction with pCDH backbone or APOBEC3C. **c.** Enrichment analyses (gProfiler) of significantly differentially expressed genes in ABM HSPCs lentivirally transduced with APOBEC3C compared to pCDH backbone-transduced controls. **d.** Heatmap of gene expression levels after unsupervised hierarchical clustering of genes enriched in gProfiler analyses and part of the KEGG Spliceosome or Reactome RNA Metabolism pathways. **e.** RNA-seq based expression levels (log counts per million) of U2AF1, ZRSR2 and SF3B2 in ABM HSPCs lentivirally transduced with APOBEC3C. **f.** Left: Number of APOBEC3-mediated C-to-U RNA editing events in close proximity, within 50 nucleotides (nt), of exon-intron boundaries normalized to the total number of edits identified. No significant differences between conditions were identified. Right: Relative number of APOBEC3-mediated single base substitutions (SBSs) within 50 nucleotides of intron-exon boundaries. Relative numbers are shown as SBSs near splice junctions and those not near splice junctions, normalized to the total number of SBSs in each category. No significant differences between conditions were observed. **g.** rMAPS motif enrichment analysis output for U2AF2 consensus binding site for exon skipping events in APOBEC3 transduced ABM HSPCs. The red line indicates motif score while the blue line indicates negative Log10 (p-value). The U2AF2 motif score is significantly enriched at a Log10 (p-value) of 1.3 or greater, equivalent to a p-value lower than 0.05. **h.** Bar plot visualizing the absolute number of SBS in ABM HSPCs upon lentiviral transduction with APOBEC3C versus pCDH. **i.** Oncoplot illustrating APOBEC3-mediated C-to-T DNA mutations following APOBEC3 lentiviral transduction of ABM HSPCs. **j.** Oncoplot illustrating APOBEC3-mediated C-to-T DNA SBS in clonal hematopoiesis (CH)-associated genes following APOBEC3 lentiviral transduction of ABM HSPCs.

**Extended Data Fig. 4:**
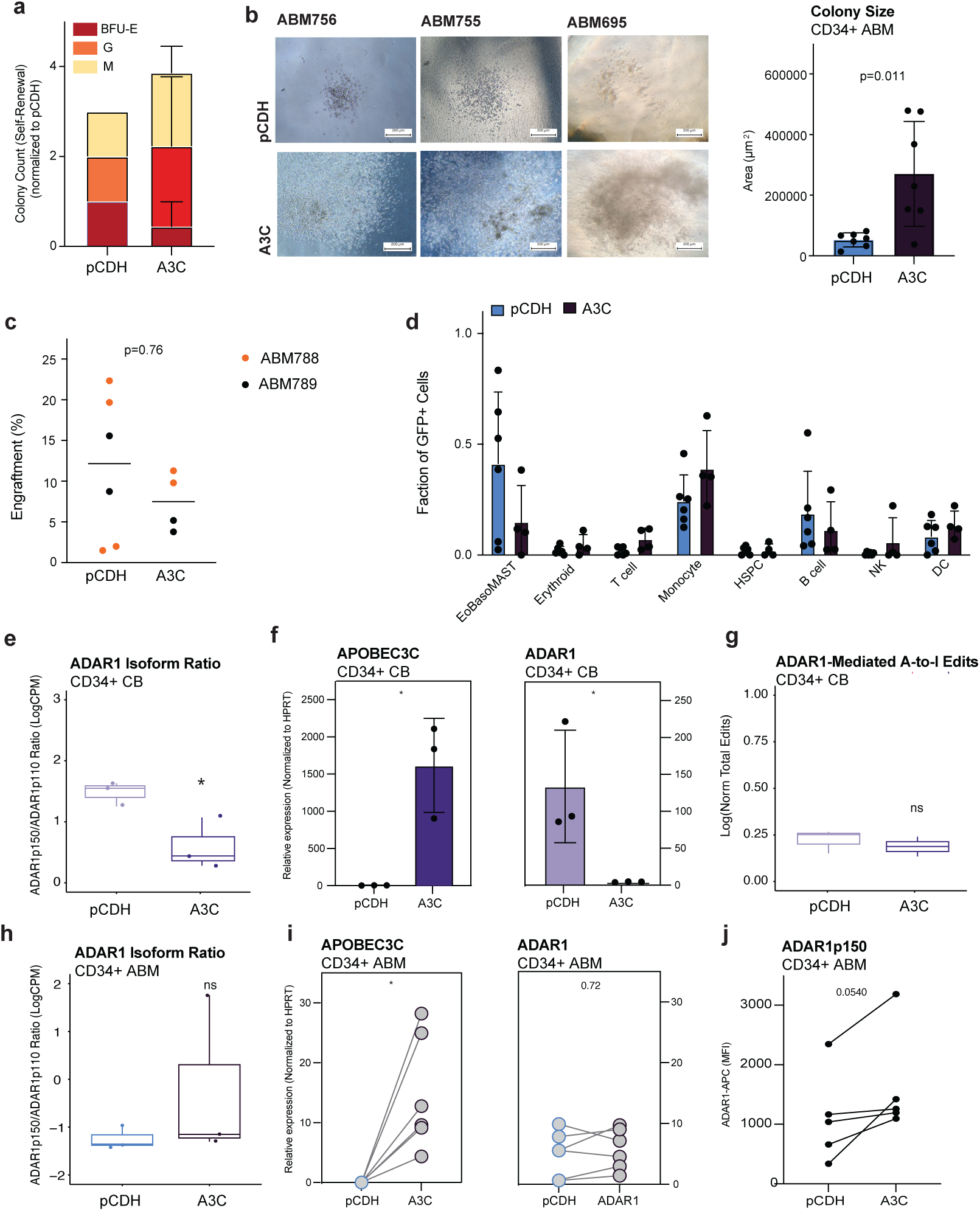
Increased hematopoietic progenitor expansion, myeloid cell frequencies and ADAR1p150 upon APOBEC3C lentiviral transduction in aged bone marrow HSPCs. **a.** Upon lentiviral transduction of ABM CD34^+^ cells with APOBEC3C or pCDH, colony counts were determined after 2 weeks of culture in methylcellulose. Counts are expressed as the percentage of colonies relative to ABM donor-matched cells transduced with pCDH. Error bars indicate SD. No statistical significance was observed. **b.** Left: Representative bright-field microscopic images of ABM CD34^+^ cells lentivirally transduced with APOBEC3C and a lentiviral pCDH backbone control. Right: Quantification of colony size. Error bars indicate SD. Statistical significance was determined using a Mann–Whitney U test. **c.** Frequencies of all human bone marrow nuclear cells upon 16 weeks of engraftment determined by single cell RNA sequencing (scRNA-seq) analyses of human bone marrow cells from mice transplanted with ABM HSPCs lentivirally transduced with pCDH or APOBEC3C. Only mice with >1% human cells detected by scRNA-seq were included in the analysis. Statistical significance was determined using a Mann–Whitney U test. **d.** Frequencies of GFP+ cells across predicted cell types based on the BoneMarrowMap reference atlas of human bone marrow hematopoiesis. Only mice with >1% human cells detected by scRNA-seq were included in the analysis. Error bars represent SD. Statistical significance was determined using a Mann–Whitney U test. No statistical significance was detected. **e.** RNA-seq-based expression levels of the ratio ADAR1 p150 to ADAR1 p110 in cord blood (CB) CD34^+^ hematopoietic stem and progenitor cells (HSPCs) lentivirally transduced with pCDH or APOBEC3C. **f.** qPCR-based expression levels of APOBEC3C and ADAR1 in cord blood (CB) CD34^+^ hematopoietic stem and progenitor cells (HSPCs) lentivirally transduced with pCDH or APOBEC3C. Error bars indicate SD. **g.** Boxplots of ADAR1-mediated A-to-I RNA edits in CB HSPCs following lentiviral transduction with pCDH backbone or APOBEC3C. **h.** RNA-seq-based expression levels of the ratio ADAR1 p150 to ADAR1 p110 in aged bone marrow (ABM) CD34^+^ HSPCs lentivirally transduced with pCDH or APOBEC3C. **i.** qPCR-based expression levels of APOBEC3C and ADAR1 in aged bone marrow (ABM) CD34^+^ HSPCs lentivirally transduced with pCDH or APOBEC3C. Error bars indicate SD. **j.** FACS analyses of ADAR1 expression in the live cell population of CD34+ selected ABM cells following lentiviral transduction with pCDH backbone or APOBEC3C.

**Extended Data Fig. 5:**
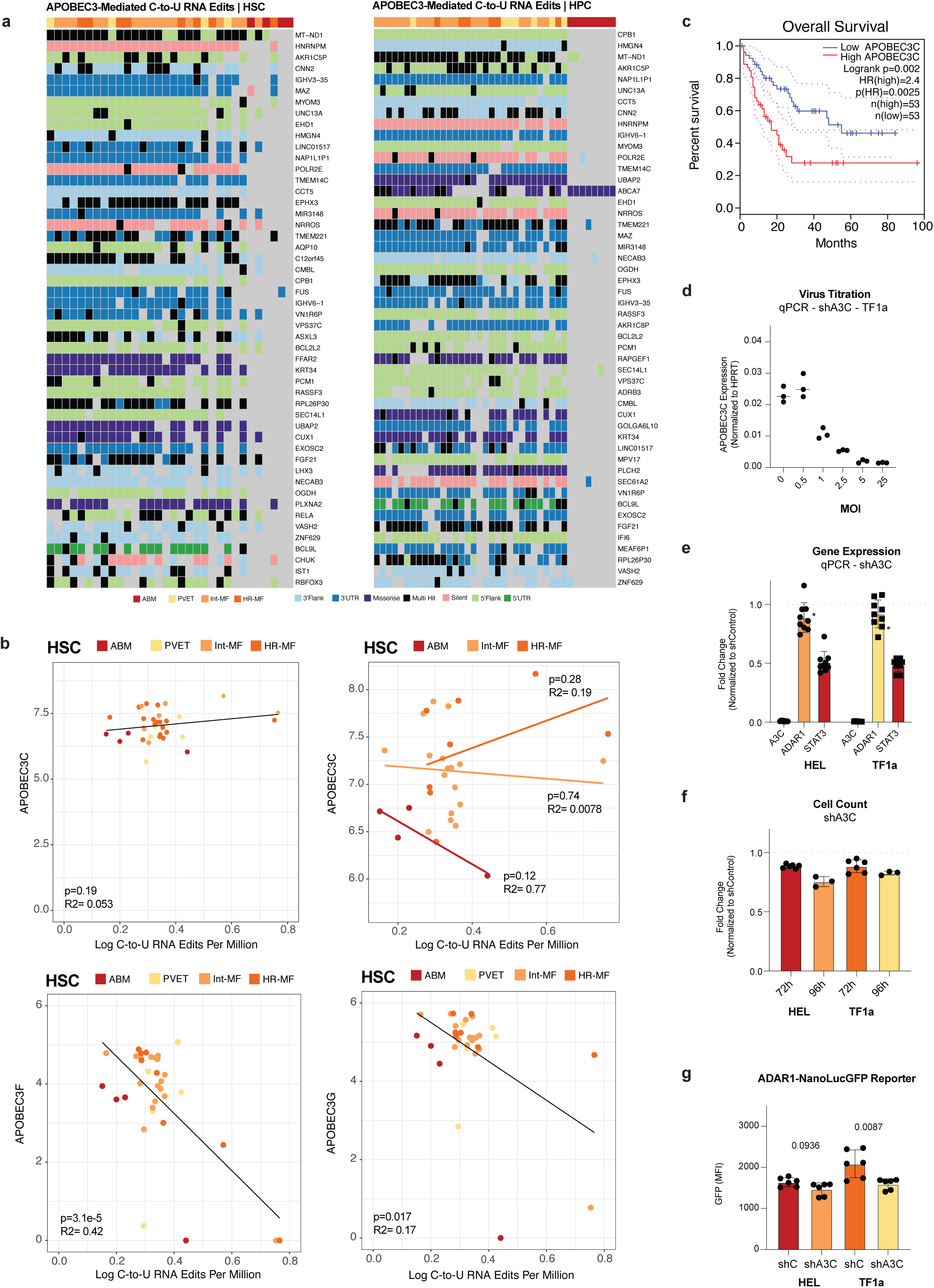
Increased C-to-U RNA editing in myeloproliferative neoplasm hematopoietic stem cells. **a.** Oncoplot illustrating the top 50 genes affected by C-to-U RNA editing in aged bone marrow (ABM), polycythemia vera (PV), essential thrombocythemia (ET), and National Comprehensive Cancer Network (NCCN) guideline-defined intermediate-risk (Int-MF) and high-risk (HR-MF) myelofibrosis. Data are shown for FACS-purified hematopoietic stem cells (HSCs; left) and hematopoietic progenitor cells (HPCs; right). **b.** Correlation of counts per million data for APOBEC3C, APOBEC3F and APOBEC3G with total C-to-U RNA edits per million reads in HSCs. and progenitors (bottom). Points are colored by disease phenotype. **c.** Kaplan-Meier plot showing overall survival (OS) of AML patients with high expression of APOBEC3C (red) or low expression of APOBEC3C (blue). **d.** qPCR based APOBEC3C expression levels upon APOBEC3C lentiviral transduction of TF1a cells at different multiplicity of infection (MOIs). **e.** Leukemia TF1a and HEL cells were lentivirally transduced with shAPOBEC3C or shControl. After 72h and 96h, cell counts were determined using FACS and are depicted as a percentage of shControl-transduced cells. Error bars indicate SD. **f.** qPCR-based expression levels of APOBEC3C, ADAR1, and STAT3 in TF1a and HEL cells 72 hours after lentiviral transduction with shAPOBEC3C, normalized to shControl. Error bars indicate SD. Significance was determined using a Mann–Whitney U test. **g.** FACS analysis of GFP mean fluorescence intensity (MFI) in the live cell population TF1a and HEL cells following sequential lentiviral transduction with our ADAR1-NanolucGFP reporter, which detects A-to-I editing, and either shAPOBEC3C or shControl. Error bars indicate SD. Significance was determined using a Mann–Whitney U test.

